# Investigating the effectiveness of cannabidiol to mitigate the adverse consequences of exposure to neonatal procedural pain

**DOI:** 10.1101/2025.02.21.639541

**Authors:** Brian Timmerman, Jennifer A. Honeycutt, Jordan Skully, Deep Patel, Waris Khan, Alan Neagu, Eshani Baez, Quinn Battagliese, Susanne Brummelte

## Abstract

Repeated painful procedures are associated with a multitude of effects on neurodevelopment in preterm infants, and current methods of neonatal pain management are unable to prevent the distress and long-term changes induced by these procedures. Cannabidiol (CBD) may be particularly effective for neonatal pain management because it reduces pain unpleasantness ratings, mitigates biological stress responses, and has minimal side effects in adults. However, there is limited research on the effects of neonatal CBD exposure. The present study investigated the efficacy of CBD treatment in mitigating behavioral responses to neonatal pain exposure. This was measured acutely through changes in ultrasonic vocalization (USV) emissions, as well as across development on long-term behavioral outcomes. We show that neonatal pain exposure decreased USV emission and increased adult anxiety-like behavior in male rats. Neonatal CBD treatment also decreased USVs in male pups but was unable to rescue the increased anxiety-like behavior in adulthood. Additionally, neonatal CBD increased baseline corticosterone levels in adult male subjects and decreased adult female body weight. More research is needed to determine whether CBD may be a safe and effective neonatal pain management medication.

**Highlights:**

- CBD reduced USV emissions on PD 2 and 3 in males, but not on later days
- Pain exposure increased adult anxiety-like behavior in males
- Neonatal CBD exposure resulted in decreased adult female body weight.
- CBD had no permanent effects on measurements of neurodevelopment or adult behavior
- CBD does not seem to mitigate effects of neonatal pain exposure

## Introduction

Infants born preterm (before 37 weeks gestational age) typically stay in the Neonatal Intensive Care Unit (NICU) after birth, where they receive medical care essential to their survival. This care necessitates exposing infants to painful procedures, such as heel pricks for blood drawing and other procedures, numerous times per day. In fact, infants in NICUs experience over a dozen painful procedures per day on average (Carbajal et al., 2008).

Exposure to a higher number of painful procedures in the NICU is associated with increased risk for a wide variety of neurobehavioral and neurodevelopmental alterations among preterm infants (as reviewed by (Grunau et al., 2013; Williams & Lascelles, 2020). For instance, infants exposed to more painful procedures in the NICU display reduced white and gray matter development at term-equivalent age (Brummelte et al., 2012), and more motor and cognitive developmental impairments during infancy (Grunau et al., 2009). At school-age, children born preterm display increased pain sensitivity (Valeri et al., 2016), increased anxiety (Ranger et al., 2014), and changes in cortical activity rhythms that are associated with impaired visuo-perceptual ability (Doesburg et al., 2013). Further, neonatal stress is observed to alter later-life hypothalamic-pituitary-adrenal (HPA) axis activity by promoting cortisol hypo-reactivity in juveniles (Brummelte et al., 2015; Chen et al., 2016; Grunau, 2013; Grunau et al., 2005; Malheiros et al., 2014) and cortisol hyper-reactivity in adults (Chen et al., 2016; (Lehmann et al., 2002; Mooney-Leber & Brummelte, 2020; Plotsky et al., 2005). Animal studies similarly indicate that in addition to prompting behavioral stress responses from pups (as measured by ultrasonic vocalization (USV) rates; (Boulanger-Bertolus et al., 2017; Takahashi et al., 1991), neonatal pain creates multidimensional changes in neurodevelopment including impaired spatial memory in juveniles (Anand et al., 2007; Chen et al., 2016; Nuseir et al., 2017; Ranger et al., 2018), changes in biological stress responses (Butkevich et al., 2023; Mooney-Leber et al., 2018; Page et al., 2005), and hyperalgesia at adulthood (Anand et al., 1999; Johnston & Walker, 2003; Nuseir et al., 2017; Page et al., 2013; van den Hoogen et al., 2021; van den Hoogen et al., 2020).

Evidence suggests that neonatal pain-associated changes in preterm infants’ neurodevelopment are driven at least in part by the secretion of glucocorticoids in response to the pain-related stressor, primarily cortisol in humans and corticosterone in rodents (CORT). CORT in turn inhibits a multitude of biological processes including immune function, inflammation, digestion, metabolism, and cell proliferation in order to prepare the body for the present stressful situation (as reviewed by (Papadimitriou & Priftis, 2009). Although this stress response can be advantageous in the short-term, elevated hypothalamic-pituitary-adrenal axis (HPA axis) activity can inhibit important neurodevelopmental processes such as neural stem/progenitor cell proliferation (Carson et al., 2016) and neuronal migration (Fukumoto et al., 2009), which could help explain the observed delayed maturation of the white and gray matter structures in infants born preterm (Brummelte et al., 2012). Effective pain management could potentially mitigate the adverse long-term effects associated with neonatal procedural pain. However, many minor procedures in the NICU (such as heel-lance or injections) are often performed without pharmacological pain management or have limited effectiveness. For instance, the primary method of pharmacological pain management for neonates during minor procedures is the non-steroidal anti-inflammatory drug acetaminophen. Although acetaminophen can reduce the need for opioids after major surgery in neonates (Walker, 2014), it is unfortunately not significantly effective for the treatment of procedural pain (as reviewed by (Ohlsson & Shah, 2020). NICUs also employ non-pharmacological methods of pain management, primarily oral sucrose and skin-to-skin care, although both methods have limitations. Oral sucrose is observed to only reduce the behavioral (but not neurological or biological) response to pain exposure (Slater et al., 2010) and is associated with developmental alterations in the dopaminergic system (McPherson & Grunau, 2014). Skin-to-skin care via physical contact with a caregiver is effective at reducing infants’ responses to procedural pain and alleviates pain-associated changes in cognitive development (Campbell-Yeo et al., 2015; Feldman et al., 2014; Johnston et al., 2008; Moore et al., 2012; Morelius et al., 2015). However, skin-to-skin care has practical limitations, as infants in NICUs are often in incubators and connected to mechanical ventilators or other medical devices that monitor their health and parents/people to provide skin-to-skin care are not always available.

It is imperative that new methods of pain management are developed for the management of frequent procedural pain in preterm infants. The endocannabinoid system (ECS), a retrograde neurotransmitter system involved in fast negative feedback at synapses, is observed to be an effective target for pain management in adult humans (as reviewed by De Vita et al., 2018) and may also be a particularly effective target for alleviating pain-related stress in neurologically immature subjects (Kwok et al., 2017; Schwaller & Fitzgerald, 2014). The ECS is involved in the regulation of HPA axis activity and nociception as well as synaptic signaling, neural progenitor/stem cell differentiation, axon migration, and synaptogenesis (as reviewed by (Corcoran et al., 2015; Fride, 2008; Harkany et al., 2008; Hill & Tasker, 2012). Modulation of the ECS could therefore potentially mitigate many of the short- and long-term consequences of neonatal pain, however, given its involvement in developmental processes, more research is needed to establish the safety of ECS manipulations during early development.

The non-intoxicating phytocannabinoid known as cannabidiol (CBD) interacts with the ECS primarily by increasing concentrations of anandamide (AEA) and has garnered a lot of attention in recent years for its ability to reduce chronic and inflammatory pain (Costa et al., 2007; De Vita et al., 2018; Gazendam et al., 2020; Hammell et al., 2016; Wong & Cairns, 2019; Xiong et al., 2012). Importantly, CBD has minimal adverse effects compared to other cannabinoids, can reduce biological stress responses, and can improve acute pain sensitivity/tolerance in adult humans (De Vita et al., 2021; De Vita et al., 2018; Gazendam et al., 2020).

While cannabinoids have exciting potential to mitigate the effects of neonatal pain-related stress, there is also good reason to be cautious about implementing them in neonatal care. Studies indicate that early-life exposure to cannabis or compounds containing tetrahydrocannabinol (THC; the primary psychoactive cannabinoid in cannabis) is associated with multiple detrimental effects (as reviewed by Grant et al., 2018; Rubino & Parolaro, 2008) including reproductive toxicity (Akingbasote et al., 2022; Carvalho et al., 2020; Corsi et al., 2021). However, the effects of THC can be drastically different from the effects of discrete CBD exposure, and exposure to stress or psychoactive substances during the prenatal period often has different effects from exposure during the neonatal period (Benoit et al., 2015; Catalani et al., 1993; McCormick et al., 2001; Modir et al., 2014). Further, most research on early-life cannabinoid exposure has focused on the consequences of cannabinoid exposure in offspring secondary to maternal cannabinoid use (as reviewed by Grant et al., 2018; Scheyer et al., 2020; Schneider, 2009; Trezza et al., 2012; Wanner et al., 2021). At present, the neurodevelopmental and reproductive consequences of direct neonatal exposure to CBD are unknown. It is therefore important that studies investigate the long-term effects of CBD itself in addition to its possible uses for neonatal pain management.

The present study investigated the potential for CBD to safely mitigate the stress and long-term developmental consequences of repeated neonatal exposure to procedural pain in a translational rodent model. We assessed pups’ ultrasonic vocalizations (USVs) in response to procedural pain, and whether USVs were reduced by pretreatment with CBD. Further, we performed a battery of developmental assessments during the neonatal period and assessed age of sexual maturity to ensure CBD had no detrimental effects on neurodevelopment or sexual maturation, respectively. At adulthood, rats underwent a series of behavioral tests for anxiety-like behavior, cognitive function, pain sensitivity, and stress responsivity, to assess if CBD could prevent the effects of neonatal pain on long-term neurobehavioral outcome.

## Methods

### Animals

Ten Sprague-Dawley female rats and four Sprague-Dawley male rats were purchased from Charles River Laboratory (Portage MI, USA) and mated to produce ten litters (two were bred a second time to produce two additional litters). All animals were housed in a 12:12-hour light: dark cycle (lights on at 7:00 AM) controlled vivarium with food and water available ad libitum.

All experiments were conducted in accordance with the National Institutes of Health Guide for the Care and Use of Laboratory Animals, and approval for all procedures was granted by the Institutional Animal Care and Use Committee of Wayne State University and can be provided upon request (Protocol #: IACUC-21-11-4217). All efforts were made to reduce the number of animals used and their suffering.

### Groups and Procedures

On postnatal day one (PD1), up to 12 pups per litter (6 males/6 females if possible) were assigned to one of six experimental groups: a) vehicle & touch, b) 5 mg/kg CBD & touch, c) 10 mg/kg CBD & touch, d) vehicle & pain, e) 5 mg/kg CBD & pain, and f) 10 mg/kg CBD & pain. If there were more than 6 pups of either sex in a litter, the surplus pups were fostered to another dam that gave birth within 48 hours and had less than 6 pups per sex. If there was no cross-foster dam available, the litter kept an uneven sex ratio and another litter got the opposite sex ratio to make up for the missing numbers (e.g. if dam A happened to have 12 males and 2 females, she kept 10 males and the 2 females (total n=12), and another dam which happened to have enough females kept 2 males and 10 females (n=12)). In any case, no more than 2 pups per sex were assigned to the same group in a litter. A total of 98 subjects (53 females / 45 males) completed the adult behavioral tests (for detailed n/group see Table 1). Any spare pups that could not be cross-fostered were culled on PD2 after CBD administration (see details below). After groups were assigned, experimenters tattooed each pup on the paw with a 26-gauge needle for identification.

**Table 1:** Number of animals per group for adult behavioral testing. One pain + 5 mg/kg male and one touch + 5 mg/kg female were euthanized between USVs and adult behavior assessment.

|  |  | Group |  |  |  |  |  |
| --- | --- | --- | --- | --- | --- | --- | --- |
|  |  | Touch & Vehicle | Pain & Vehicle | Touch & 5 mg/kg | Pain & 5 mg/kg | Touch & 10 mg/kg | Pain & 10 mg/kg |
| Sex | Female | 10 | 9 | 8 | 8 | 9 | 9 |
|  | Male | 8 | 7 | 7 | 8 | 8 | 7 |

### Drug Preparation and Administration

The first two litters were gavaged during PD1-4, which unfortunately resulted in some PD1 pups needing to be euthanized and thus the administration window was moved to PD2-5 for subsequent litters (n=8) to ensure feasibility and survival of the pups. For the gavaging procedure, pups were first weighed and separated from their dams into a holding cage on top of a heating pad. Pups were separated for at least 20 minutes before intraoral drug administration to ensure they had sufficient gastric capacity. Intraoral drug administration of CBD or vehicle was performed via oral gavage with a 22-gauge metal gavage needle. Briefly, an investigator held the pup by the scruff and skin of the upper torso so as to tilt the pup’s head back. The investigator then inserted the gavage needle into the mouth at a 45° angle before rotating the gavage needle around its tip to a 90° angle and lowering the needle into the pup’s esophagus. The investigator then slowly administered the drug solution (maximum 10 µl per second) before carefully removing the gavage needle. Another investigator then monitored gavaged pups for complications before returning them to their home cage.

CBD was administered as Epidiolex (purchased from GW Pharmaceuticals/Jazz Pharmaceuticals) which contains CBD, dehydrated alcohol (7.9% w/v), strawberry flavor, and sucralose in solution with sesame seed oil. Because the volume of Epidiolex that needed to be administered daily to each pup was quite small (0.05 µl/g or 0.10 µl/g for a pup in the 5 mg/kg group or 10 mg/kg group, respectively), additional sesame oil was added to the Epidiolex solution to increase volumes to 10 µl/g and make administration easier. Vehicle solution consisted of sesame oil, as the Epidiolex placebo was not available for purchase. Pups received 10 µl/g of either vehicle (sesame oil), 5 mg/kg CBD, or 10 mg/kg CBD via oral gavage. Epidiolex and vehicle solutions were stored at room temperature per manufacturer instructions for Epidiolex.

### Pain/Touch and Ultrasonic Vocalization Recording

Seventy-five minutes after drug administration, pups were transferred from their home cage into a holding cage on top of a heating pad. One at a time, pups received either procedural pain in the form of a 26-gauge needle inserted into a paw, or control touch in the form of a paintbrush touched against both sides of a paw as previously described (Mooney-Leber et al., 2018). Needle pokes and touches were alternated across paws such that each paw was used no more than once per day. Pain/touch exposure was performed four times per day on PD2-5 at 75, 150, 255, and 330 minutes after drug administration. Immediately after pain/touch exposure each pup was placed in a sound attenuating cubicle (Med Associates, Inc., ENV-022M) and ultrasonic vocalizations (USVs) were measured via a USV microphone (Avisoft Bioacoustics, CM24/CMPA) for 120 seconds. Recordings were scored for the number and total duration of vocalizations between 30 – 85 kHz via Avisoft SASLab Pro (v5.3.01, Avisoft Bioacoustics). After 120 seconds of USV recording the pup was returned to the holding cage. To avoid pups being separated from their dam for too long, only half the litter was in a holding cage at a time. After the first half of the litter completed the session, they were returned to their home cage and the second half of the litter was placed in the holding cage. The order in which subjects were tested was pseudo-randomized. For the sake of feasibility, USVs were not recorded during the third poke/touch exposure and pups were instead immediately returned to their holding cage after stimulus exposure (see Figure 1 for an experimental timeline).

**Figure 1:**
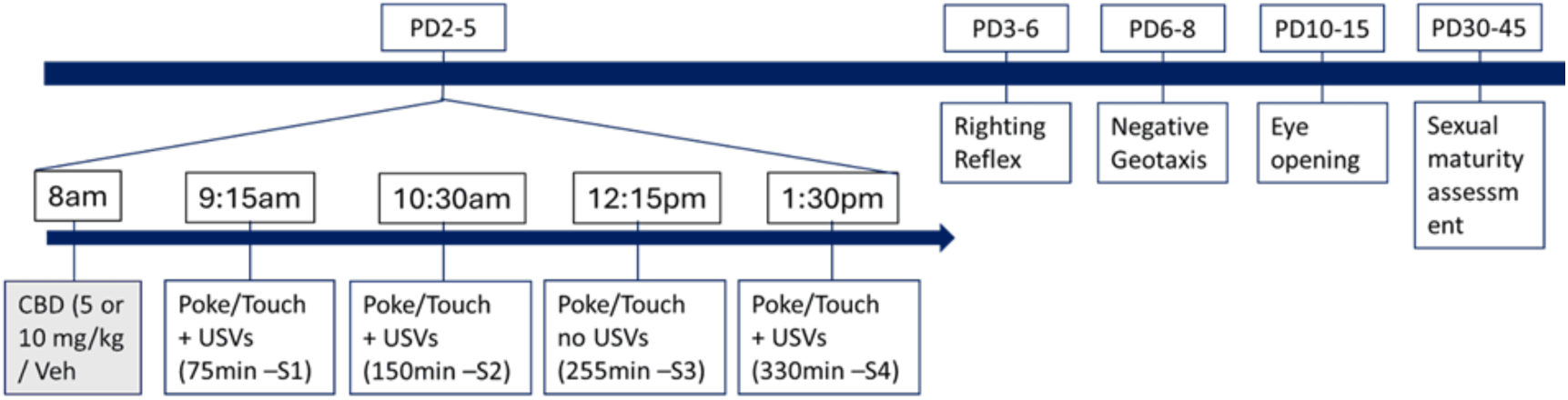
Developmental Timeline. Cannabidiol (CBD) or Vehicle (Veh) was administered once in the morning from Postnatal day (PD) 2-5, followed by Pain/touch exposure at 75, 150, 255, and 330 minutes after drug administration. USVs were recorded immediately after Session (S) 1,2 and 4. A battery of assessments for developmental milestones were performed throughout the neonatal and juvenile period.

#### USV Network training

A Faster region-based convolutional neural network (Faster-RCNN) was trained in DeepSqueak using over 2,800 training images. These images included a mix of calls from rats in the present study as well as those from rats across different developmental age groups, which were collected in-house. Files selected for training displayed diverse categories of calls (Wright et al., 2010) in order to increase model effectiveness (Gong et al., 2019). USVs from initial training audio files were manually selected to train the network. Calls were first identified using call characteristics that have been previously classified (Wright et al., 2010; Shair, 2018). Selections were then made in DeepSqueak that contained the entire call. Network training images were collected using parameters defined by (Coffey et al., 2019-11-06) for short-duration vocalizations. The faster-RCNN was trained three times using different images to reduce call detection errors. Following training, calls within all audio files were detected using the network and then manually checked and corrected when necessary by an experimenter blind to condition.

#### USV Call correction, parameters, and analysis

All files included ultrasonic calls that had harmonics which mirrored the shape of the actual call, though at a higher frequency (Coffey et al., 2019; Grimsley et al., 2011). When harmonics appeared on the spectrogram, only the call at the lowest frequency was manually accepted as valid (Fonseca et al., 2021). Each detection file was manually corrected by an experimenter, blind to condition, to include all valid calls between 30 kHz and 85 kHz. The score, call length, principal frequency, high frequency, and low frequency were extracted for each detected call within a given file. These data were evaluated to determine average values for each session and the frequency of valid calls was analyzed for each day and group.

### Plasma CBD Concentration

On PD2 all spare pups (i.e. if a dam had more than 12 pups and the 2 extra litters, n=22 (4M/18F)) were weighed and separated from their dam alongside littermates. During drug administration spare pups received either 5 mg/kg or 10 mg/kg of CBD. These spare pups were then sacrificed either 75 minutes or 150 minutes later (to line up with the time of the 1st and 2nd USV sessions). Trunk blood was collected at sacrifice in EDTA vials and samples were kept at 4° C until processed. Samples were spun down at 2500 rpm for 15 minutes for plasma extraction. Plasma was then kept at -20° C until it was analyzed for CBD concentration via liquid chromatography-mass spectrometry by the Lumigen Instrument center service at Wayne State University.

### Neonatal Developmental Battery

A battery of developmental measures selected from standard behavioral teratology screening protocols (Adams & Jones, 1984) were used to assess if neonatal CBD exposure influences subjects’ neurodevelopment (see descriptions of tests below; see Figure 1 for an overview of the developmental timeline). Body weight (g) was assessed at the beginning of the day on PD1-5 and then periodically until weaning (PD8, PD12, PD16 & PD21). Surface righting reflex was assessed on PD3-6. An investigator placed a rat pup on their back on a flat surface and then measured the latency (in seconds) for the rat pup to turn fully upright. Negative geotaxis was assessed daily on PD6-8. An investigator placed a pup on a 35° incline facing downwards and then measured the latency (up to 60 seconds) for the pup to turn to face upwards (turn 180°), after which the pup was returned to their home cage. Eye opening was assessed daily starting on PD10 until both eyes were open for each pup. Investigators recorded the age at which each eye first opened for each subject. Pups were weaned on PD21 and group-housed with same sex siblings. Once subjects reached the cage weight limits, they were pair-housed with a same sex sibling or housed in groups of 3 if necessary (once one ‘cage’ reached the weight/size limits to require pair-housing, all animals in the litter switched to pair housing to keep conditions similar for all subjects). All animals were checked daily from PD 30-45 for age of reaching sexual maturity defined as vaginal opening for females and testis descent for males.

### Adult Behavioral Assessment

Subjects performed a series of behavioral tests at young adulthood (starting at ∼ PD60). Animals were allowed to recover for at least 24 hours between each test (see Figure 2 for a timeline of adult behavioral assessments) and handled for at least 3 days prior to the first behavioral test.

**Figure 2:**
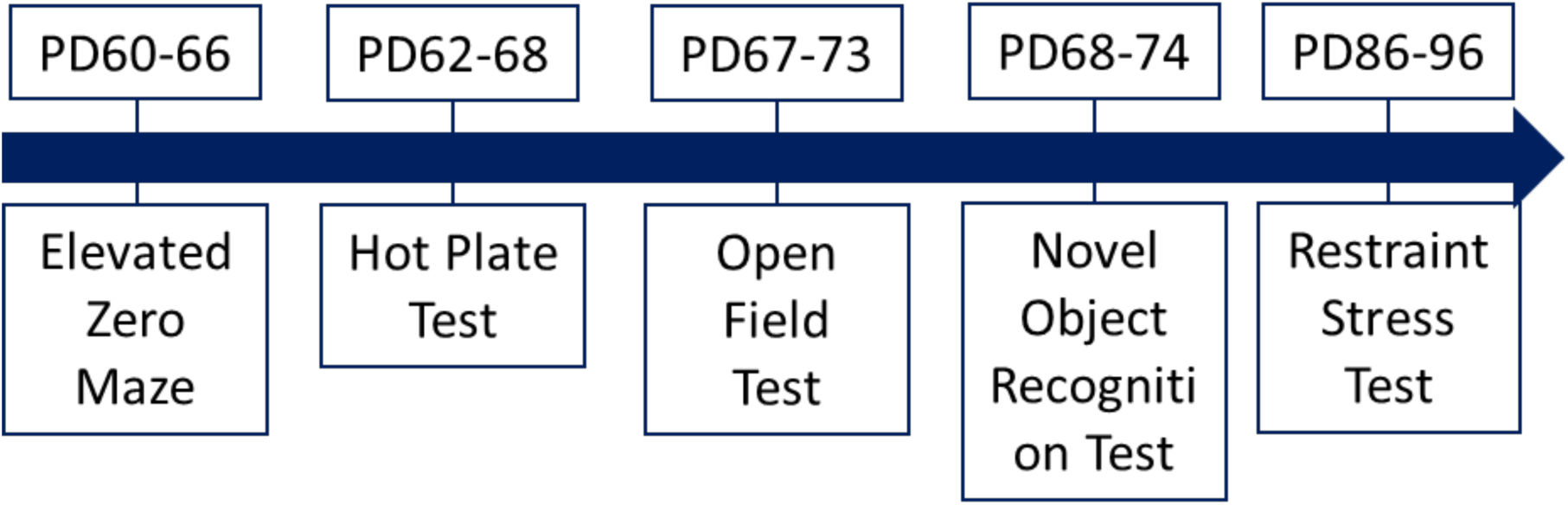
Timeline of Adult Behavioral Tests. Each test was only performed once per animal, but animals’ age at time of testing varied slightly due to varying birth dates.

### Estrus Cycling

Adult female animals were lavaged to determine their estrous cycle phase on the day prior to each behavioral test as well as the day of the test to assess any effect of estrous stage on behavior. Briefly, three to four drops of water were administered into the vagina via an eyedropper and then immediately reabsorbed. From the eyedropper, samples were placed on a glass slide and stained with Cresyl Violet. Once dry, samples were analyzed under a microscope and estrous phase was scored by an experimenter based on the types of cells observed (McLean et al., 2012). To control for possible influences of the lavage procedure on behavior, male subjects underwent a “mock lavage” (i.e., a gentle stroke of the genitals with a paintbrush) any time female littermates were lavaged.

### Elevated Zero-Maze (EZM)

Subjects performed the EZM test once during ∼PD60-66 to assess anxiety-like behavior. The maze (Med Associates, Inc) consisted of a ring-shaped platform (86.4 cm in height and 61 cm in diameter) with 2 closed sections (8.1 cm walls) on opposite sides of the maze. An experimenter placed the subject onto the EZM in front of the entrance to a closed section and then immediately left the room. Experimenters videotaped the subject’s behavior and analyzed the amount of time spent in open/closed areas and the distance traveled in open/closed areas via behavioral tracking software (Noldus Ethovision XT 16). After five minutes in the maze, the subject was removed and returned to their home cage. Experimenters then removed any fecal boil and cleaned the maze with Accel TB before the next subject began the test.

### Hot Plate Test

Subjects performed the hot plate test once between PD62-68 to assess pain thresholds via a rodent Analgesiometer (San Diego Instruments). The day before a subject was tested, experimenters placed the rat on the analgesiometer while it was turned off to familiarize subjects with the environment. On the day of testing, an experimenter placed the rats in the testing room with their lab coat less than 6 inches away from the cages for 30 minutes to allow them to habituate to the room and olfactory stimuli from the experimenter. Afterwards, subjects were individually placed on the analgesiometer set to 50° C. The experimenter immediately removed the subject from the hot plate upon the initiation of paw-licking, or within 60 seconds if the subject did not react. The experimenter then recorded the subject’s latency to paw lick and cleaned the hot plate with Accel TB before running the next subject.

### Open Field Test (OFT)

Subjects performed the OFT once between PD67-73 to assess anxiety-like behavior. Adult subjects were placed in a square arena (80 cm x 80 cm x 36 cm) in low-light conditions and allowed to explore for 10 minutes while their behavior was recorded via a video camera (acA1300, Basler, Hamburg Germany). Videos were scored via Ethovision XT 17 (Noldus, Wageningen, Netherlands) for distance traveled, time spent in the inner 50% of the arena, latency to enter the inner area, and number of entries into the inner area. After removing a subject from the arena, experimenters removed any fecal boil and cleaned the arenas with Accel TB before the next subject began the test.

### Novel Object Recognition

Subjects performed the novel object recognition test the day after the OFT (i.e., between PD 68-74) to assess recognition memory. Animals were placed in the same arena from the OFT but with two identical objects positioned equidistant from each other and the walls of the arena. Each subject was allowed to explore the arena and objects for 10 minutes before they were returned to their cage. Subjects that did not investigate the objects for at least 10 seconds during this phase were excluded from further analysis. One hour after returning to their cage, subjects were placed in the arena again with one of the previous objects replaced by a new and unfamiliar object. Animals were allowed to explore the arena and objects for five minutes before they were returned to their cage. During both sessions, subjects’ behavior was recorded by a video camera (acaA1300, Basler, Wageningen, Netherlands). Recordings were scored by behavioral tracking software (Noldus Ethovision XT 17) for distance traveled, time spent investigating each object, and the frequency of object visits. Novel object preference was calculated as the time spent investigating the novel object divided by the total time spent investigating either object (i.e., novel object time / (novel object time + familiar object time)).

### Restraint Stress Test and Serum Corticosterone

Subjects underwent restraint stress testing once between PD86-96 to assess stress responsiveness and HPA axis function. A baseline blood sample was collected via saphenous vein blood draw (t0) within 3 minutes of an experimenter touching the subject’s cage. The subject was then immediately placed in a plastic restraint stress tube (Plas labs, model 553-BSSR for females & 554-BSSR for males) for 30 minutes. Once the subject was removed, experimenters collected a second blood sample (t30) before returning the subject to their home cage. One hour later, a third blood sample was collected from the subject (t90). No more than 7 ml/kg was collected from each subject.

Blood samples collected during the Restraint Stress Test were kept at 4° C and allowed to clot overnight. The next day, samples were spun down at 8000g for 10 minutes for serum extraction. Serum was then kept at -20° C until it was analyzed for corticosterone concentrations using a standard CORT EIA kit (catalog # K014-H1; Arbor assays, Ann Arbor MI) per manufacturer’s recommendations. All samples were run in duplicates and the intra assay coefficient was less than < 0.10.

### Euthanasia & Adult Weight

Subjects were sacrificed around eight days after they underwent restraint stress testing. All animals were weighed, anaesthetized with pentobarbital and then transcardially perfused and brains were extracted for future use.

#### Statistics

All data were analyzed via Generalized Estimating Equation (GEE) modeling with exchangeable working correlation matrices and gamma-log model structure. GEE modeling was chosen instead of repeated measures ANOVA to reduce the influence of clustering (i.e., violations of the assumption of the homogeneity of variance) from litter effects. Because logarithmic models cannot be calculated if a value equals 0, all dependent variables that contained null values had all values increased by 1 exclusively for statistical tests and estimated marginal means calculation. This was done for USVs, EZM entries into the open arm, and OFT entries into the center of the arena. Factors in all analyses included sex (male, female), pain condition (pain, touch), and CBD condition (vehicle, 5 mg/kg, 10 mg/kg). For all repeated measures (neonatal weight, USVs, righting reflex, negative geotaxis, and restraint stress CORT), each measurement point was included in their respective models as within-subjects factors. To reduce type 1 error rate, all non-significant interaction effects involving within-subject factors were removed via backward elimination starting with the highest order interactions (e.g., if day, pain, and drug were factors in an analysis and there were no significant interactions by day, then the day by drug by pain interaction was removed, followed by day by drug and day by pain but not drug by pain). Further, significant effects from tests that included within-subjects factors were followed-up with pairwise comparisons using Bonferroni correction. All other significant effects were followed-up with pairwise comparisons using least significant differences. All effects were considered significant at p < 0.05. Extreme outliers were defined as any values more than 3 standard deviations from the mean. Assessments of subjects in adulthood were also followed-up by tests of exclusively female subjects, controlling for estrus cycle phase (proestrus/estrus vs diestrus/metestrus) as covariates. All statistical analyses were performed with IBM SPSS 29.

## Results

### Ultrasonic Vocalizations

Equipment issues during the first few weeks of USV recording necessitated the removal of subjects in litters A, B, C, and D from USV analysis (thereby also removing USV values from the two litters that received pain and/or CBD on PD1-4 instead of PD2-5). To prevent further reductions in sample sizes, the values for any extreme univariate outliers (n= 3) from the remaining pups were changed to match the next-closest value recorded from a subject in the same group at the same postnatal age and session number (e.g., an extremely high value recorded from a male touch + vehicle subject on PD3 session 2 would be replaced with the next-highest value recorded on PD3 session 2 from another male touch + vehicle subject (Tabachnick & Fidell, 2013).

Analysis of USVs found a significant main effect of sex (Wald χ2 = 14.83, p < 0.001), with male subjects producing more vocalizations than female subjects on average (see Figure 3A). As a result, all subsequent USV analysis was performed separately by sex.

**Figure 3:**
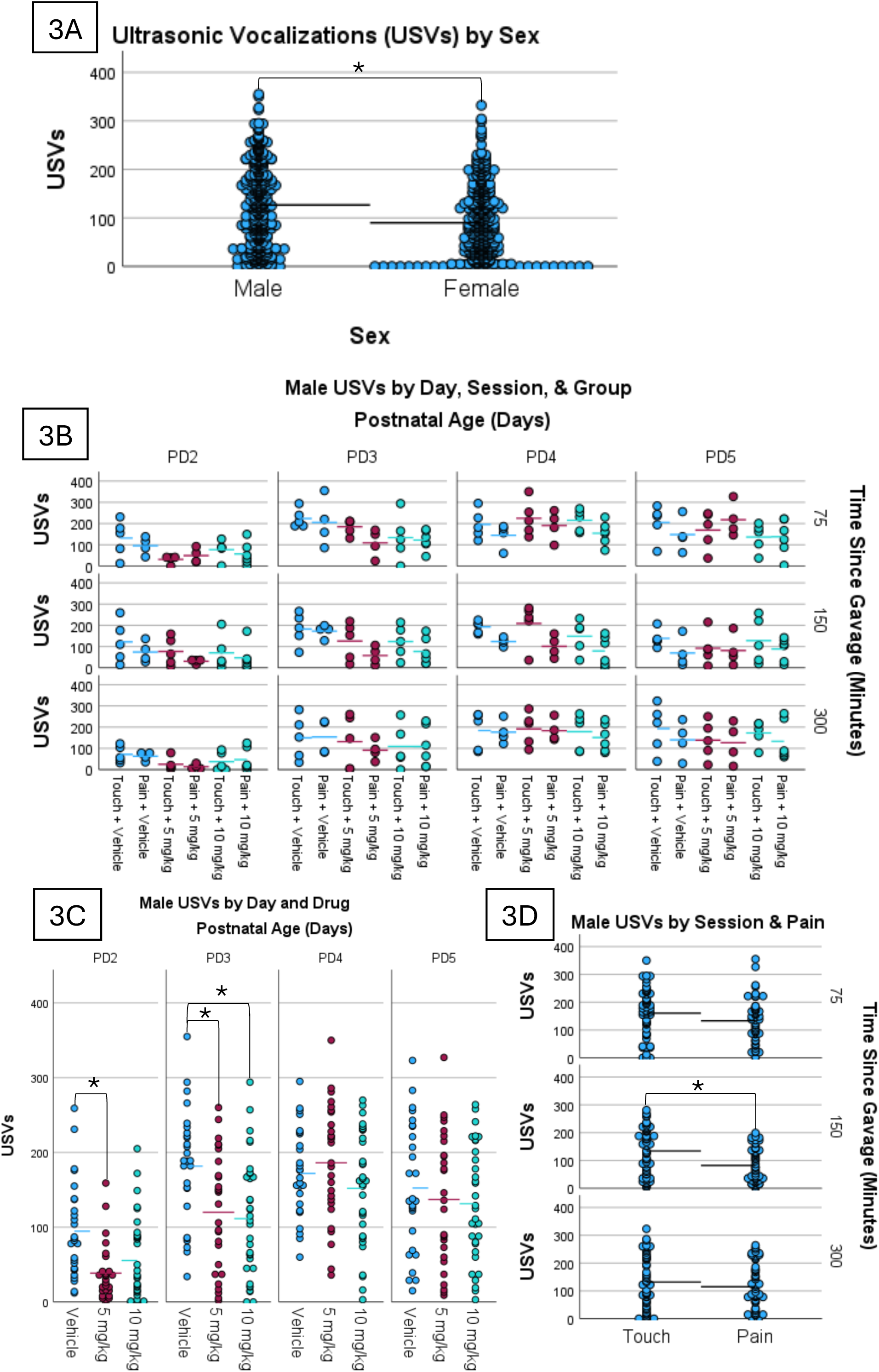
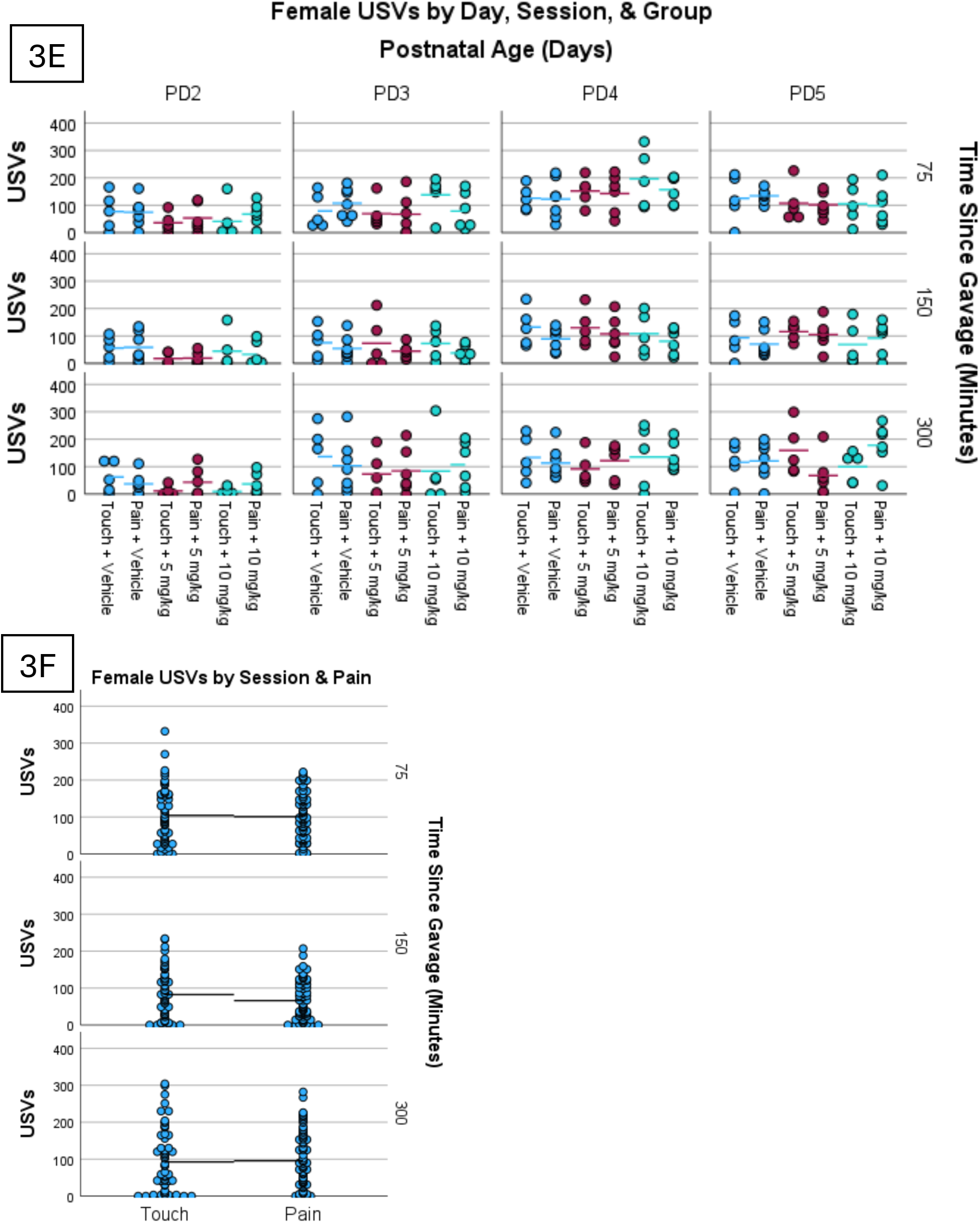
USVs. The data presented in this Figure is from the same subjects but separated by sex and collapsed across different variables to highlight the significant effects. All USVs collected are visualized separately for males (A) and females (E). Male subjects produced significantly more vocalizations than female subjects (A). Among males, vehicle subjects produced significantly more vocalizations compared to 5 mg/kg CBD subjects on PD2 as well as compared to 5 mg/kg and 10 mg/kg CBD subjects on PD3 (C). Among males, pain subjects produced significantly more vocalizations than touch subjects during session 2 but sessions 1 or 3 (D). Although there was a significant session by pain interaction among females, no significant pairwise comparisons survived Bonferroni correction (F). * p < 0.05.

USVs in males were significantly impacted by day (Wald χ2 = 128.09, p < 0.001), session (Wald χ2 = 23.29, p < 0.001), drug (Wald χ2 = 19.40, p < 0.001), pain (Wald χ2 = 7.38, p < 0.007) as well as days and drug (Wald χ2 = 47.32, p < 0.001) and Session and pain (Wald χ2 = 12.15, p = 0.002) interactions (Figure 3B). After Bonferroni corrections, pairwise comparisons revealed that CBD-exposed male pups produced fewer USVs than vehicle pups on PD 2 (5 mg/kg group, p < 0.001) and on PD 3 (5 and 10 mg/kg group, p’s < 0.009; see Figure 3C). Further, male pain pups produced fewer USVs compared to male touch pups during session 2 (i.e., 150 min after gavage; p = 0.001; see Figure 3D).

In females, USVs were also significantly impacted by day (Wald χ2 = 84.64, p < 0.001), and session (Wald χ2 = 57.69, p < 0.001), and session by pain (Wald χ2 = 7.72, p = 0.02, all data displayed in Fig, 3E), but there were no main effects of drug or pain and follow-up pairwise comparisons for the session by pain interaction revealed no pain effect at any session (only differences between sessions that varied slightly between pain and touch animals (see Figure 3F).

### Plasma CBD Concentration

Analysis of plasma CBD concentrations found a significant main effect of time (Wald χ2 = 12.08, p < 0.001) with CBD concentrations being significantly higher in subjects 150 minutes after administration than 75 minutes after administration (see Table 2). There were no significant effects for dose (p’s > 0.05).

**Table 2:** Plasma CBD concentration. Assessment of CBD concentration in plasma at 75 minutes and 150 minutes after oral administration of 5 mg/kg or 10 mg/kg CBD found no significant difference between drug conditions, but did find a significant increase between 75 and 150 minutes (mean ± S.E.M, all p’s > 0.05).

| Time (min) | Dose | Mean ± S.E.M. | Group size |
| --- | --- | --- | --- |
| 75 | 5 mg/kg | 73.3 ± 20.3 | 4 |
| | 10 mg/kg | 69.1 ± 5.9 | 4 |
| 150 * | 5 mg/kg | 120 ± 27.1 | 8 |
| | 10 mg/kg | 168.7 ± 28.8 | 7 |

### Surface Righting and Neonatal Geotaxis

Litters A and B (n = 12) were removed from this analysis because their drug and pain exposure occurred on different postnatal days than other litters (PD1-4 instead of PD2-5). This decreased the number of days that pain and/or CBD exposure overlapped with righting reflex assessment in litters A and B, and therefore likely altered their righting reflex performance.

The GEE test for righting reflex found significant main effects for day (Wald χ2 = 114.55, p < 0.001) and drug (Wald χ2 = 10.0, p = 0.007), as well as significant interaction effects for day by pain (Wald χ2 = 10.64, p = 0.014), day by drug (Wald χ2 = 21.70, p = 0.001), and day by sex by drug (Wald χ2 = 17.37, p = 0.008; see Figure 4A). Pairwise comparisons for day by drug by sex found that 10 mg/kg CBD males were slower to right themselves than touch males on PD4, while 5 mg/kg CBD females were slower than vehicle females on PD5 and also slower than both vehicle and 10 mg/kg CBD females on PD6 (see Figure 4A) Pairwise comparisons for the day by pain interaction found that righting latencies in the touch group significantly decreased PD4-5 and PD5-6, but pain group latencies significantly decreased only between PD5-6 (see Figure 4B).

**Figure 4:**
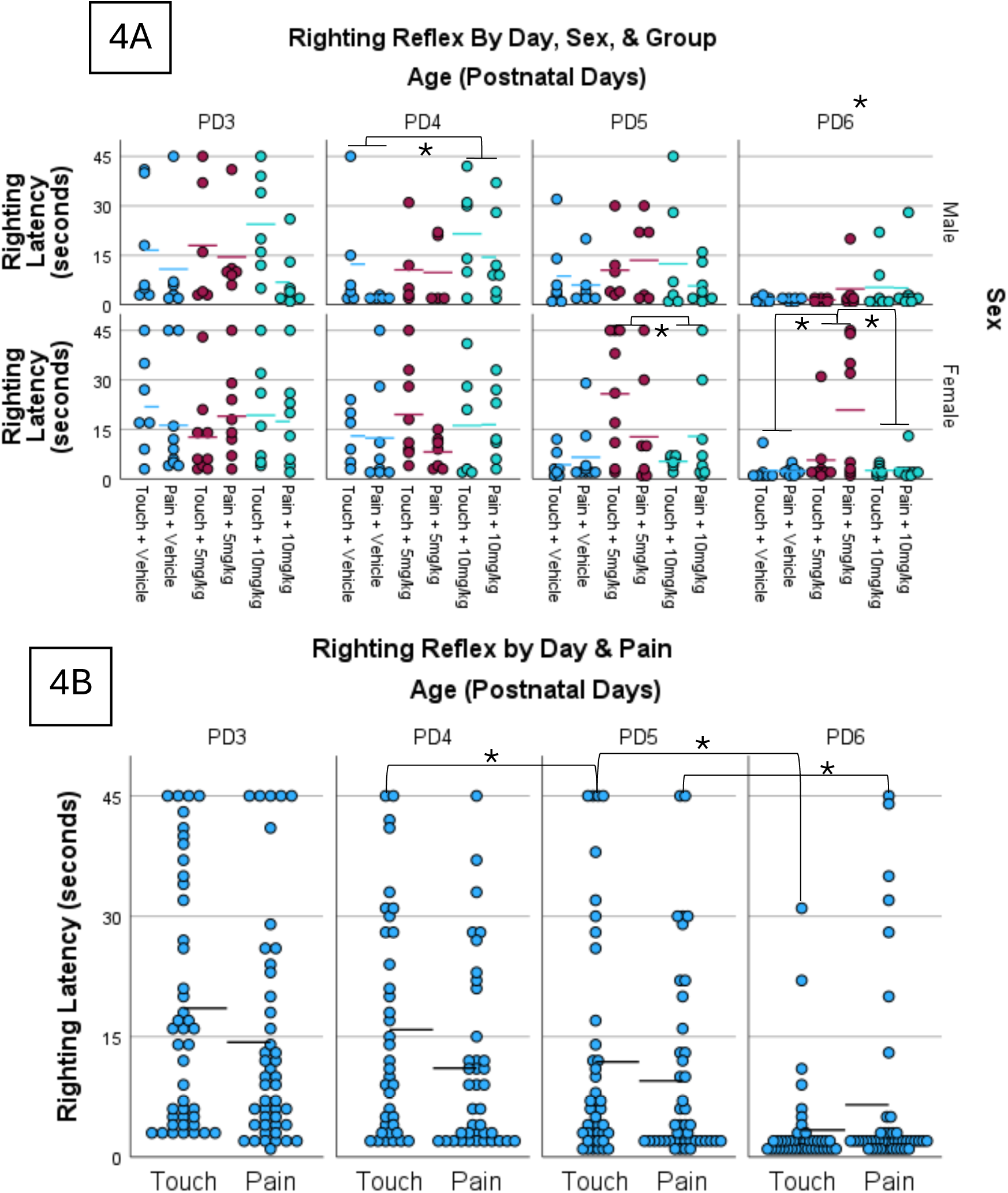
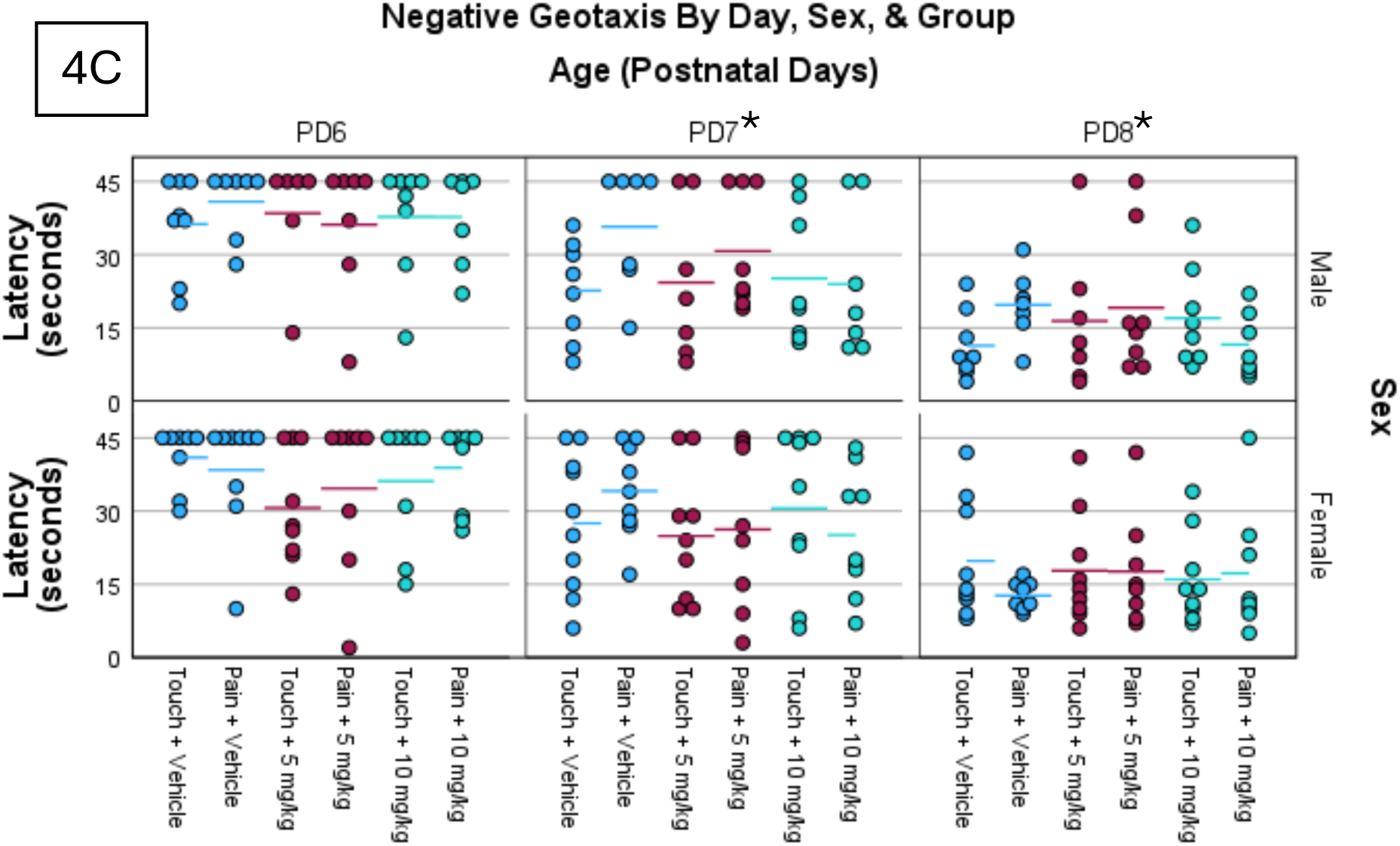
Neuromotor reflex assessment. Overall surface righting performance did not significantly change between PD3-5 but significantly decreased between PD5-6 (A). Ten mg/kg CBD impaired male righting reflex performance on PD4, while 5 mg/kg impaired female performance on PD5 and PD6 (A). Touch subjects improved significantly between PD4-6, while pain subjects only improved significantly between PD5-6 (B). Negative geotaxis latency decreased between PD6-8 but did not significantly differ between groups (C). * p < 0.05

The GEE test for negative geotaxis found a significant main effect of day (Wald χ2 = 164.6, p < 0.001), driven by subjects’ latency decreasing over PD6-9 (see Figure 4D). No other significant effects or significant interactions were found.

### Neonatal Body Weight

Assessment of subjects’ weights on PD1-5 and PD8, PD12, PD16 and PD21 found a significant effect of age (Wald χ2 = 68451.7, p < 0.001) and sex (Wald χ2 = 18.1, p < 0.001; see Table 3). There was also a significant age by sex interaction (Wald χ2 = 25.73, p = 0.001), with male subjects weighing more than female subjects on PD1-16 but not PD21. No other significant effects were found for neonatal weight (all other p’s > 0.05).

**Table 3:** Developmental Milestones. There was no significant difference between treatment groups or pain vs touch for age at eye opening, age to reach sexual maturity (Vaginal opening/ testis drop) or body weight until the end of weaning. There was a significant effect of sex and sex by day interaction for body weight starting on PD 1 until PD16 (all p’s<0.05). All data are displayed as means ± S.E.M.

| Sex | CBD/Veh | Pain/Touch | Age at Eye Opening [d] | Age at Sexual Maturity [d] | Weight PD1 [g] | Weight PD4 [g] | Weight PD8 [g] | Weight PD12 [g] | Weight PD16 [g] | Weight PD21 [g] |
| --- | --- | --- | --- | --- | --- | --- | --- | --- | --- | --- |
| Male | Vehicle | Touch | 13.9 ± 0.2 | 39.6 ± 0.8 | 7.7 ± 0.3 | 11.5 ± 0.5 | 19.3 ± 0.9 | 28.4 ± 1.2 | 37.6 ± 1.6 | 52.7 ± 2.2 |
|  |  | Pain | 13.9 ± 0.2 | 40.0 ± 0.5 | 7.1 ± 0.1 | 10.5 ± 0.2 | 17.6 ± 0.4 | 26.0 ± 0.5 | 34.4 ± 0.6 | 48.2 ± 1.0 |
|  | 5 mg/kg | Touch | 14.1 ± 0.2 | 39.6 ± 0.8 | 7.1 ± 0.2 | 10.6 ± 0.3 | 17.8 ± 0.5 | 26.3 ± 0.7 | 34.7 ± 0.9 | 48.7 ± 1.4 |
|  |  | Pain | 14.1 ± 0.3 | 40.4 ± 0.9 | 7.0 ± 0.2 | 10.4 ± 0.3 | 17.5 ± 0.4 | 25.8 ± 0.7 | 34.1 ± 0.8 | 47.8 ± 1.3 |
|  | 10 mg/kg | Touch | 14.1 ± 0.3 | 40.3 ± 1.0 | 7.6 ± 0.3 | 11.3 ± 0.5 | 18.9 ± 0.9 | 27.9 ± 1.2 | 36.9 ± 1.6 | 51.7 ± 2.3 |
|  |  | Pain | 13.9 ± 0.3 | 39.5 ± 0.4 | 7.6 ± 0.3 | 11.3 ± 0.4 | 19.0 ± 0.7 | 28.0 ± 1.1 | 36.9 ± 1.4 | 51.8 ± 2.0 |
| Female * | Vehicle | Touch | 13.9 ± 0.2 | 38.8 ± 0.3 | 6.5 ± 0.1 | 9.8 ± 0.2 | 16.8 ± 0.3 | 24.9 ± 0.4 | 33.2 ± 0.6 | 46.6 ± 0.9 |
|  |  | Pain | 14.0 ± 0.2 | 38.2 ± 0.1 | 6.9 ± 0.3 | 10.4 ± 0.5 | 17.9 ± 0.8 | 26.5 ± 1.2 | 35.4 ± 1.5 | 49.7 ± 2.1 |
|  | 5 mg/kg | Touch | 13.7 ± 0.3 | 38.3 ± 0.2 | 6.7 ± 0.1 | 10.1 ± 0.2 | 17.3 ± 0.3 | 25.7 ± 0.4 | 34.4 ± 0.5 | 48.3 ± 0.8 |
|  |  | Pain | 14.1 ± 0.2 | 39.3 ± 0.4 | 6.6 ± 0.2 | 9.9 ± 0.2 | 17.1 ± 0.4 | 25.3 ± 0.6 | 33.8 ± 0.7 | 47.4 ± 1.1 |
|  | 10 mg/kg | Touch | 14.1 ± 0.3 | 39.0 ± 0.4 | 6.7 ± 0.1 | 10.1 ± 0.2 | 17.3 ± 0.2 | 25.7 ± 0.3 | 34.3 ± 0.4 | 48.2 ± 0.7 |
|  |  | Pain | 13.9 ±<br>0.2 | 38.3 ± 0.2 | 6.6 ±<br>0.1 | 9.9 ±<br>0.2 | 17.0 ±<br>0.4 | 25.2 ±<br>0.6 | 33.7 ±<br>0.7 | 47.3 ±<br>1.0 |

### Eye opening and Age at Sexual Maturity

GEE analysis of postnatal age at eye opening found no significant effects for sex, pain, drug, or any interactions of these variables (all p’s > 0.05; see Table 3).

GEE analysis of postnatal age at sexual maturity found a significant main effect of sex (Wald χ2 = 14.26, p < 0.001) with males reaching sexual maturity at a significantly later age than females (see Table 3;). There were no significant effects of pain and/or CBD.

### Adult Behavioral Tests

#### Hot Plate Test

GEE analysis of paw-lick latency during the hot plate test found a significant effect of sex (Wald χ2 = 8.34, p = 0.004) driven by female rats having significantly longer paw-lick latency than male rats (see Table 4), but no other significant effects were found. A follow-up test of only female subjects with estrous stage as a factor found that proestrus combined with estrus (Wald χ2 = 0.02, p = 0.88) had no significant effect on paw-lick latency.

**Table 4:** There was no significant difference in the latency to withdraw paws from the hot plate apparatus (Sec ± S.E.M.; all p’s > 0.05) or in the discrimination index for the Novel Object Task ( Index, ± S.E.M, all p’s > 0.05).

| Sex | CBD/Veh | Pain /Touch | Hot Plate Paw withdrawal latency [s] | Novel Object Discrimination index |
| --- | --- | --- | --- | --- |
| Male | Vehicle | Touch | 18.2 ± 2.4 | 0.47 ± 0.03 |
|  |  | Pain | 21.2 ± 3.1 | 0.56 ± 0.06 |
|  | 5 mg/kg | Touch | 17.2 ± 2.8 | 0.51 ± 0.02 |
|  |  | Pain | 19.3 ± 2.9 | 0.56 ± 0.05 |
|  | 10 mg/kg | Touch | 21.0 ± 3.2 | 0.52 ± 0.05 |
|  |  | Pain | 18.4 ± 2.6 | 0.63 ± 0.03 |
| Female | Vehicle | Touch | 16.1 ± 2.0 | 0.59 ± 0.04 |
|  |  | Pain | 13.9 ± 1.0 | 0.48 ± 0.08 |
|  | 5 mg/kg | Touch | 13.9 ± 1.7 | 0.51 ± 0.08 |
|  |  | Pain | 13.6 ± 2.4 | 0.54 ± 0.07 |
|  | 10 mg/kg | Touch | 11.6 ± 1.4 | 0.47 ± 0.06 |
|  |  | Pain | 18.2 ± 2.7 | 0.68 ± 0.08 |

#### Elevated Zero Maze

GEE analysis found significant main effects of sex on time spent in the open arms of the EZM (Wald χ2 = 4.25, p = 0.039), for the number of entries into open arms (Wald χ2 = 5.60, p = 0.018) and distance traveled in the EZM (Wald χ2 = 6.09, p = 0.014). Female subjects moved significantly farther, and entered open arms more frequently, and spent more time in the open arms than male subjects (see Figure 5A, 5C, & 5D). Thus, all subsequent analysis was performed separately by sex.

**Figure 5:**
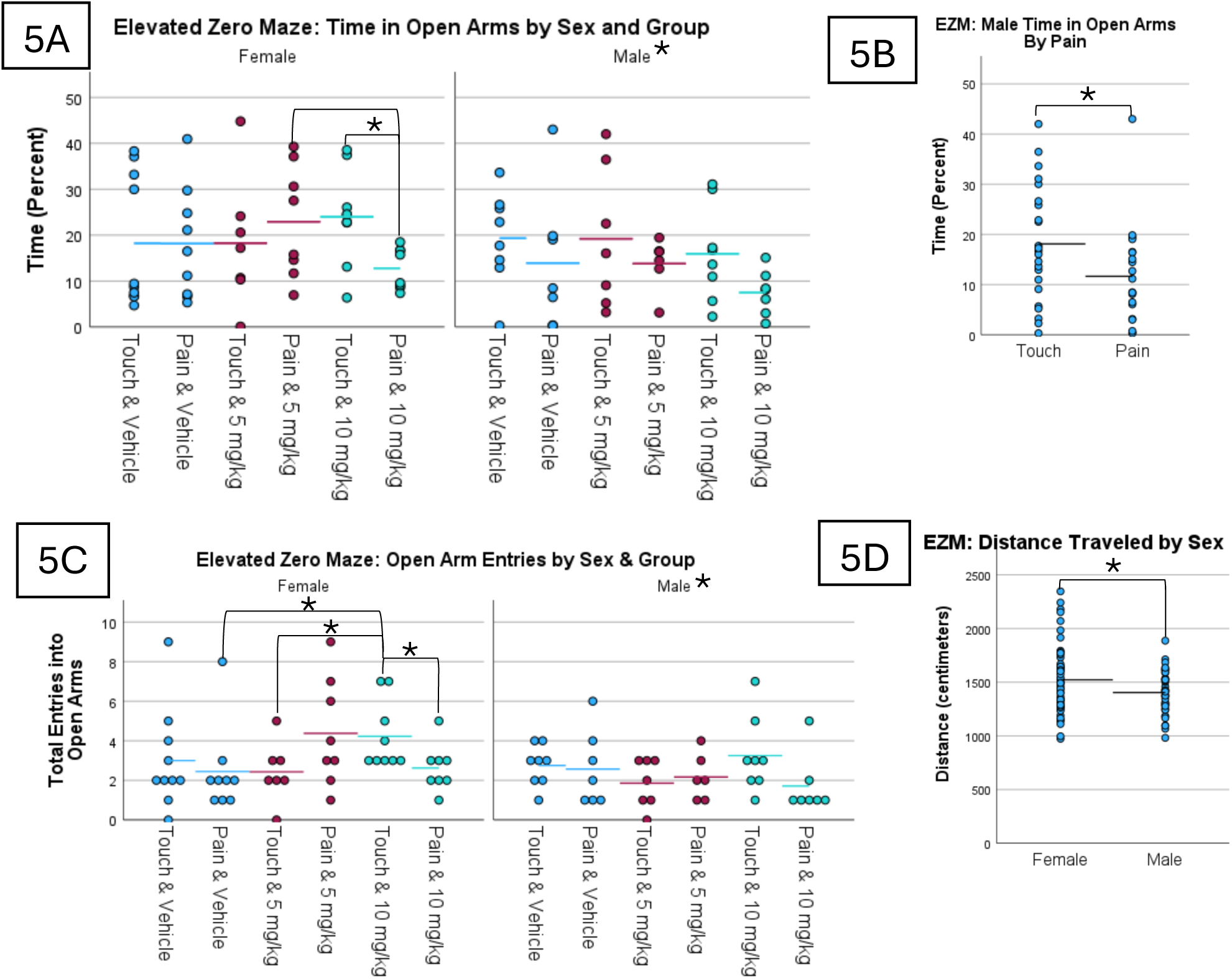
Elevated Zero Maze Performance. Among females, pain & 10 mg/kg subjects spent significantly less time in open arms compared to touch & 10 m/kg subjects and pain & 5 mg/kg subjects. Male subjects spent significantly less time in open arms of the EZM than female subjects (A). Among males, pain subjects also spent significantly less time in open arms than touch subjects (B). Females also entered open arms more frequently than males, and among female subjects the touch & 10 mg/kg group entered open arms significantly more frequently than the pain & vehicle, touch & 5 mg/kg, and pain & 10 mg/kg group (C). Female subjects also traveled significantly farther than males (D). * p < 0.05

For females, GEE analysis revealed a significant interaction effect of pain and drug for the percent of time spent in the open arms (Wald χ2 = 6.90, p = 0.032) as well as the number of open arm entries (Wald χ2 = 8.95, p = 0.011). Pairwise comparison revealed that females in the 10 mg/kg pain group spent less time in the open arm compared to the 5 mg/kg pain group (p = 0.019) and compared to the 10mg touch group (p = 0.001; see Figure 5A). The same pattern was observed for number of entries into the open arms, with females in the pain & 10 mg/kg group also making significantly fewer entries into the open arms compared to touch & 10 mg/kg females (p = 0.017) and (non-significantly) fewer entries compared to the pain & 5 mg/kg group (p = 0.075; see Figure 5C). Further, the touch & 10 mg/kg females made more entries into open arms compared to the touch & 5 mg/kg females (p = 0.018). There were no group effects for total distance travelled among females, and proestrus combined with estrus did not have a significant effect on time spent in the open arms (Wald χ2 = 0.003, p = 0.96), number of entries (Wald χ2 = 0.902, p = 0.342), or distance traveled in the EZM (Wald χ2 = 0.28, p = 0.60).

For males, GEE analysis revealed that pain subjects spent significantly less time in the open arms of the maze than touch subjects (Wald χ2 = 5.37, p = 0.02; see Figure 5B), but there were no effects for number of entries into the open arm or distance travelled (all p’s > 0.05).

#### Open Field Test

Open field test data from two subjects were excluded from analysis because their trials were interrupted. GEE analysis found significant mains effect of sex for distance traveled in the arena (Wald χ2 = 47.80, p < 0.001) and the number of entries into the center (i.e., inner 50% of the arena; Wald χ2 = 10.34, p = 0.001) and thus follow-up GEEs were performed separately by sex.

Among female subjects, there were no group or interaction effects for the percent of time spent, latency to enter or number of entries into the center of the open field or the total distance travelled (all p’s> 0.05, Figure 6A, 6C, & 6E), and there was no significant effect of proestrus or proestrus and estrus combined on any of these measures either (all p’s> 0.05).

**Figure 6:**
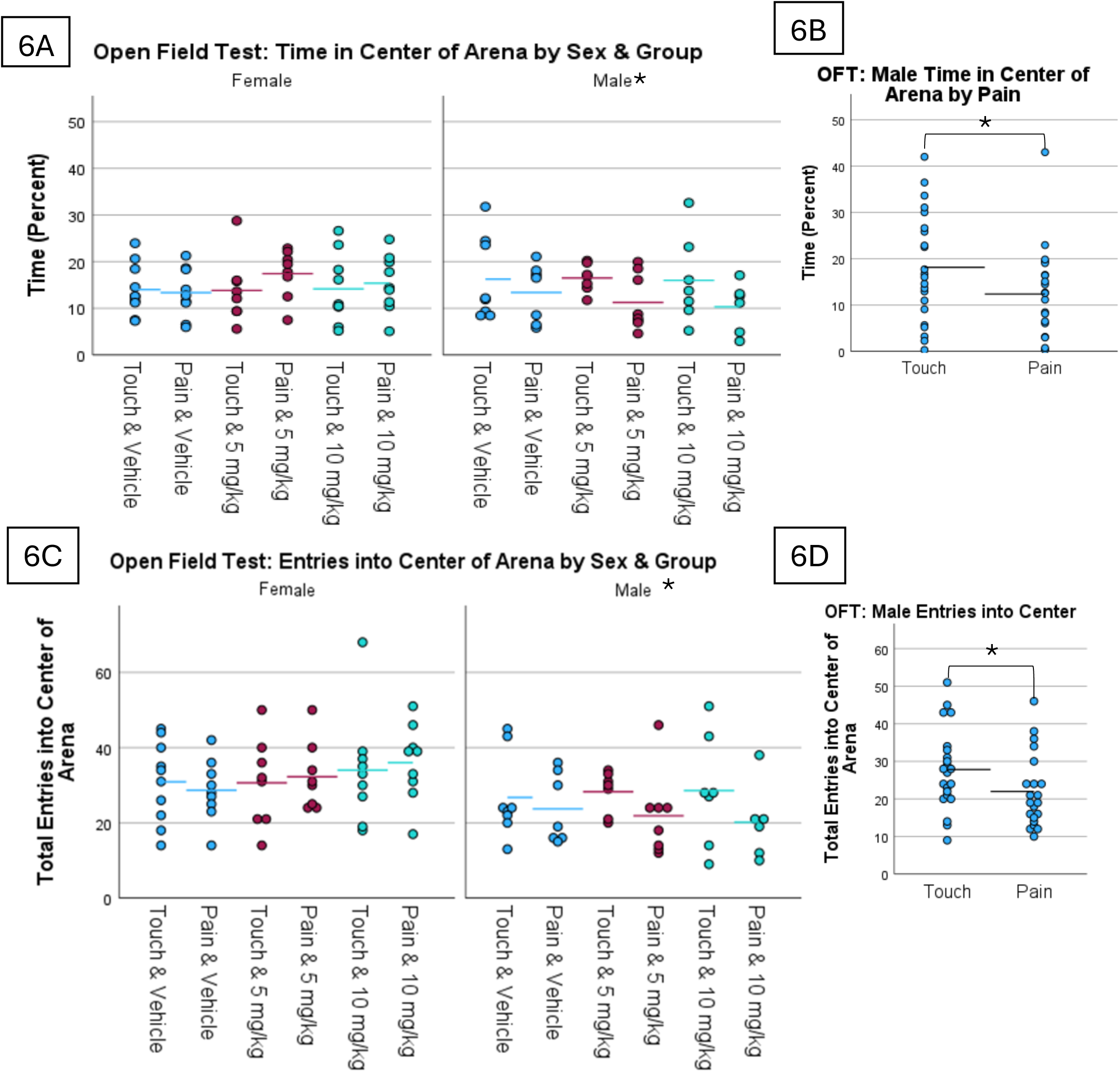
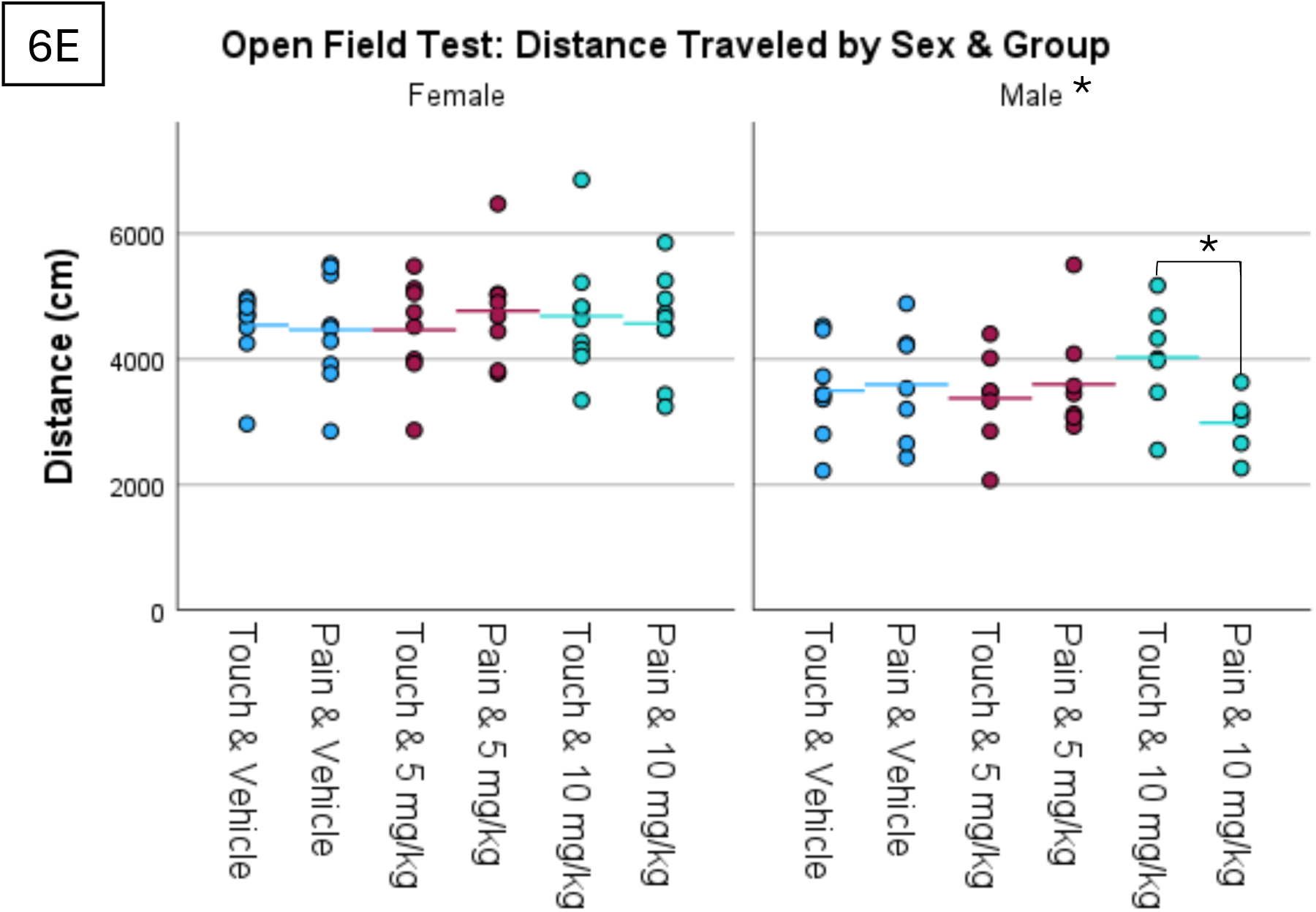
Open Field Test Performance. Compared to female subjects, male subjects spent significantly less time in the center of the Open Field Arena (A) and entered the center fewer times (C). Among male subjects, the touch group entered the center of the arena fewer times and spent less time there compared to the pain group (B, D). Male subjects also traveled significantly less than female subjects, and among males the pain & 10 mg/kg group traveled significantly less than the touch & 10 mg/kg group (E). *\* p < 0.05*

For males, there was a significant effect of pain for time in the center (Wald χ2 = 5.96, p = 0.015) and number of entries into center (Wald χ2 = 3.85, p = 0.050) with male pain subjects spending significantly less time in the center of the arena and making fewer entries than male touch subjects. There was also a pain by drug interaction effect for total distance traveled (Wald χ2 = 7.91, p = 0.019, Figure 6B and 6D) and the pairwise comparison revealed that this was driven by the 10 mg/kg pain males travelling less than the 10 mg/kg touch males (p=0.003). Further, the latency to enter the center revealed a significant main effect of drug (Wald χ2 = 6.09, p = 0.048), but the follow-up analysis revealed no specific significance between the groups and the mean latency for all three groups to enter the center was within 1-2 sec of being placed in the arena, so this potential difference may not be very meaningful (data not shown).

#### Novel Object Recognition Test

Any subjects that did not spend at least 10 seconds total investigating objects during the familiarization or novel object trials were excluded from analysis (n=3). GEE analysis of distance moved during the novel object trial found a significant effect of sex (Wald χ2 = 78.27, p < 0.001), with female subjects moving significantly more than male subjects (, but there were no significant effects for the discrimination index for any of the groups or for females only after accounting for the estrus cycle (all p’s <0.05, see Table 4).

#### Restraint Stress Corticosterone Response

Male and female subjects were assessed separately because of inherent sex differences in CORT concentrations that were also clear in the current study (main effect of sex: Wald χ2 = 255.941, p < 0.001).

The test of male subjects found a significant main effect of time (Wald χ2 = 962.1, p < 0.001) as well as a significant effect of time by drug (Wald χ2 = 12.33, p = 0.015) driven by 10 mg/kg CBD males having higher baseline CORT than vehicle males (see Figure 7A) as well as a non-significant effect of pain (Wald χ2 = 3.72, p = 0.054), with pain exposed animals seeming to have slightly lower corticosterone levels.

**Figure 7:**
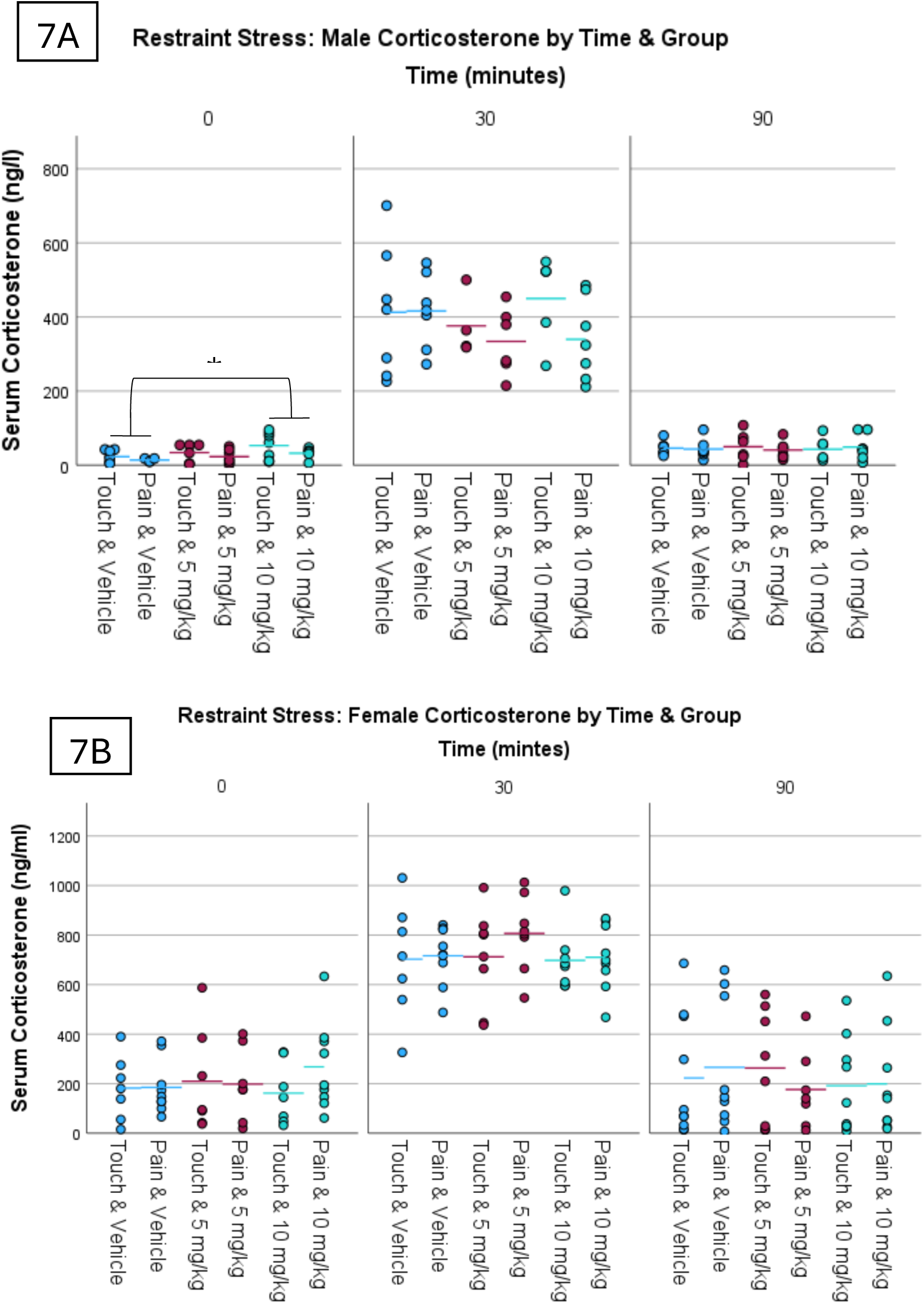
Corticosterone Response to Restraint Stress from Males (A) and Females (B). Male 10 mg/kg CBD subjects had significantly lower CORT levels compared to male vehicle subjects at baseline (A). Female subjects displayed no significant overall effects for CBD or pain (B). * p < 0.05

Assessment of female CORT concentrations found a significant main effect of time (Wald χ2 = 161.26, p < 0.001; see Figure 7B) with estrus cycle phase being a significant covariate in that analysis (Wald χ2 = 4.79, p = 0.029), but no other significant effects.

#### Adult Body Weight

Male and female subjects were assessed separately due to a clear sex effect in adult body weight (Wald χ2 = 2675.26, p < 0.001). No significant effects for weight were found among adult male subjects (see Figure 8). However, adult females displayed a significant main effect of drug (Wald χ2 = 7.09, p = 0.029) and a significant drug by pain interaction effect (Wald χ2 = 16.09, p < 0.001). Follow up analysis revealed that vehicle dams weighed more than 10 mg/kg CBD dams (p = 0.013), while female pain + vehicle subjects weighed significantly more than female touch + 5 mg/kg subjects (p = 0.003) and female pain plus 10 mg/kg subjects (p < 0.001). Further, female pain + 5 mg/kg subjects also weighed more than female pain + 10 mg/kg subjects (p = 0.003; see Figure 8). There was no significant effect of the estrus cycle phase on adult female body weight (Wald χ2 = 2.22, p = 0.14).

**Figure 8:**
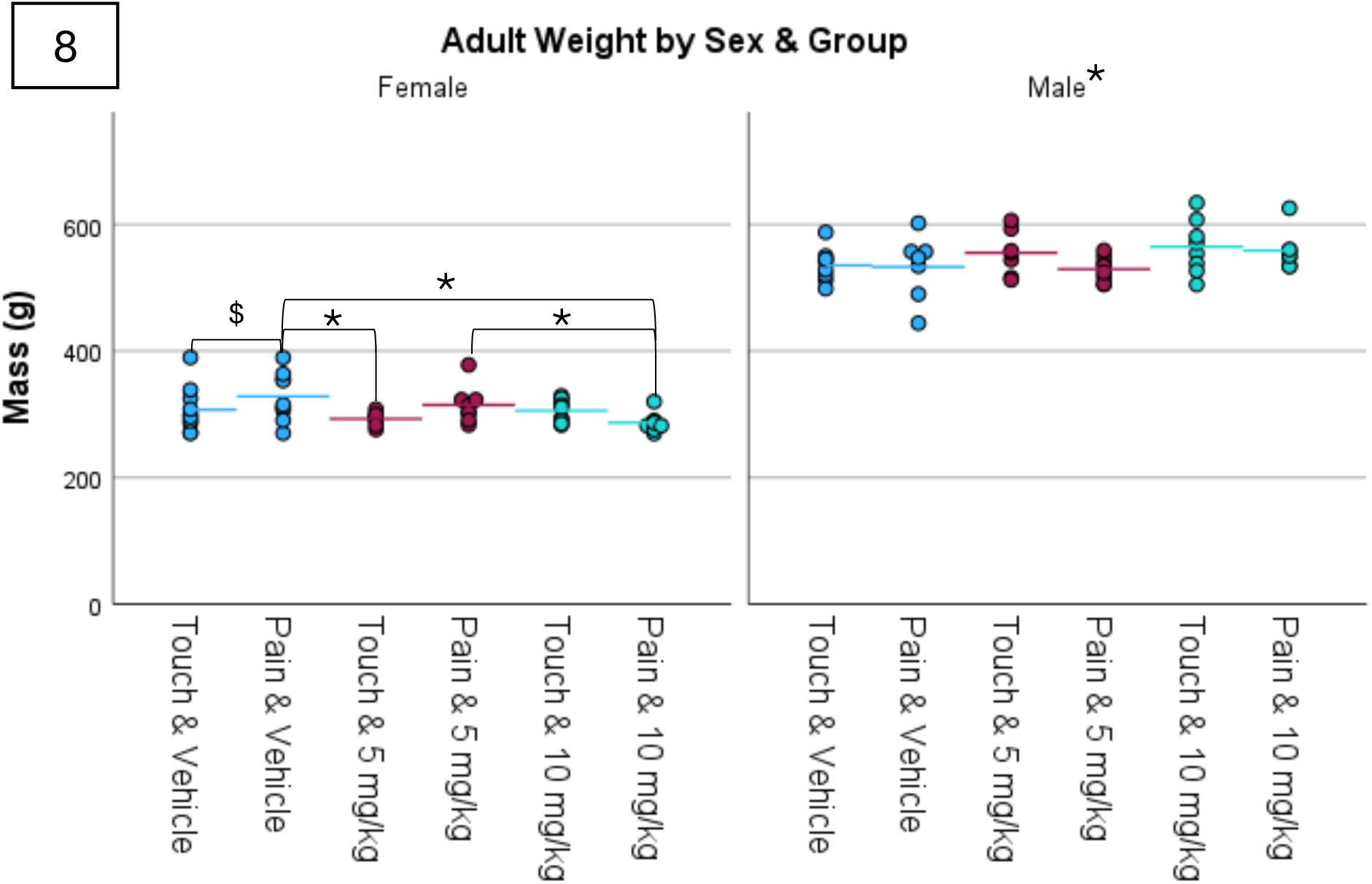
Adult Weights. Males weighed significantly more than females, but neither pain nor drug condition had a significant effect on males’ adult weights. Among female subjects, the pain plus vehicle group weighed significantly more than the touch plus 5 mg/kg group and pain plus 10 mg/kg group, and the pain plus 5 mg/kg group weighed significantly more than the 10 mg/kg group. * p < 0.05, $ p < 0.05 compared to 10 mg/kg subjects.

## Discussion

Our results revealed that both neonatal pain and CBD treatment decreased USVs, although CBD’s effect on USVs ceased by day 4. Contrary to our hypothesis, we did not see any pain by CBD interactions, suggesting that CBD was not able to mitigate any of the adverse effects of neonatal pain exposure in our rodent model. CBD itself seemed to transiently slow maturation of neuromotor performance, with CBD pups displaying slower self-righting performance compared to vehicle controls but no difference in subsequent assessments of negative geotaxis. Additionally, male 10 mg/kg CBD subjects had slightly higher baseline corticosterone levels at adulthood and CBD appeared to decrease the adult body weight of female subjects. On the other hand, neonatal pain increased anxiety-like behavior especially in adult males, independent of any CBD treatment.

### Complicated Relationship Between Neonatal Stress/CBD

#### Exposure and USVs

To our knowledge, the present study is the first to demonstrate that neonatal CBD administration influences USV emission in rat pups. Both doses of daily CBD administration reduced neonatal vocalization rates in male pups on postnatal day 3, and 5 mg/kg CBD also reduced neonatal vocalizations on postnatal day 2, while there was no CBD effect in female pups and no effects were observed on Days 4 or 5. These findings indicate that neonatal CBD administration may be having relieving effects, but these are only observed in male pups, and decrease after repeated administration. Our findings are in line with prior studies that investigated gestational and/or postnatal exposure to other drugs that increase endocannabinoid signaling, such as whole cannabis extract or CB receptor agonists, and also found decreased neonatal USVs (Manduca et al., 2012; Manduca et al., 2020). While adult rats are capable of emitting USVs reflective of both appetitive and aversive affective states (Wöhr & Schwarting 2013), pups tend to emit USVs during periods of distress, specifically when separated from their mothers (Winslow and Insel, 1991). The aversive nature of these calls is supported by previous research showing anxiolytic drugs attenuate pup USV emissions, whereas anxiogenic drugs increase these calls (Insel et al., 1986, Rainer et al., 2018). Based on these results, the decrease in USV count in male rats administered 5mg and 10mg doses suggested that neonatal CBD administration has an anxiety-reducing effect.

It is possible that the sex-specific effects of neonatal CBD administration on USV count may be reflective of inherent differences between male and female USV emission. Previous research has highlighted that from PN2 to PN13, males emitted a greater number of USVs than females (Naito & Tonneau, 1987; Bowers et al., 2014). Additionally, the calls emitted by female pups were shorter in duration (Naito & Tonneau, 1987). Interestingly, a sex effect is also reported in human studies of neonatal vocalization with neonatal males being more vocal than females (Oller et al., 2020).

Given that CBD has multiple mechanisms that may be relevant to stress and pain management, including interactions with the ECS, dopamine receptors, serotonin 1A receptors, and adenosine uptake (as reviewed by Mlost et al., 2020), it is not that surprising that CBD may influence pain-related vocalizations (Blazevic et al., 2017; Ise & Ohta, 2009). Though we did not find pain by CBD interaction effects, we found that procedural pain reduced vocalizations, with USVs in session 2 (i.e. 150 minutes after gavaging CBD or vehicle) driving this effect. Further, our follow-up analyses showed that this effect was only significant in male pups, although females showed a similar, yet non-significant, pattern. This pain-related reduction in USVs seems to be in contrast with the theory that decreased USVs indicate decreased anxiety/distress (Simola, 2015; Winslow & Insel, 1991). However, the explanation likely lies in the ethological function of neonatal USVS. Neonatal USVs promote retrieval and care by dams (Boulanger-Bertolus, J. et al., 2017), but USV production is suppressed when pups experience predator stress as a part of an adaptive response to avoid predation (Branchi et al., 2001; Takahashi, 1992; Wiedenmayer et al., 2003; Zuluaga et al., 2023). Therefore, pups in the present study may have decreased USV production after pain exposure because they perceived it as a threatening or fear-inducing stimulus and their dams were not present in the chamber. Consistent with this, Takahashi et al (1990; 1991) found that footshocks beginning immediately after pups are placed into a chamber decreased their USV production, while Boulanger-Bertolus et al. (2017) found increased USVs in neonatal pups (PD 12-15) exposed to footshocks after habituation to a chamber or in response to rough maternal handling, stimuli that are likely painful but not fear-inducing.

Taken together, both, CBD exposure and pain each reduced USVs at particular time points and especially in males, but contrary to our hypothesis we did not observe a specific interaction between CBD and pain that would suggest a pain-relieving effect of CBD in pain-exposed pups. Future research investigating neonatal stress responses via USVs should consider the environmental context of the stress/pain exposure to ensure that it is not fear-inducing and try to further tease apart the ethological meaning/relevance of USVs in varying scenarios.

#### Plasma CBD Concentration

CBD concentrations in plasma samples were higher at 150-minutes than 75-minutes after administration but did not significantly differ by dosage. Interpretation of this particular finding is somewhat limited by our small sample size for this measure, but the data is relatively similar to what we have previously found in adult rats (levels peak around 2 hours after administration and then start to decrease; unpublished data) and suggest that the absorption of CBD after oral administration may occur at a relatively constant but slow rate. Given the timing of our pain-exposure sessions in this experiment, the plasma concentration data suggests that pups should have experienced the greatest acute CBD effect during their 2nd pain session. Interestingly, this is the session where we see the strongest pain effect on USVs, however there was no interaction effect for pain and CBD for that session, suggesting that the pain effect is not driven by the higher levels of CBD during that time point.

#### Neurodevelopment

Neonatal administration of CBD significantly increased righting reflex latency on PD5-6 but not PD3-4. Notably, CBD only differed from vehicle when vehicle subjects’ performances began to improve, which suggests that CBD specifically prevented improvement in neuromotor performance. Importantly, the observed effect in our study cannot be attributed to acute drug effect as the testing was performed in the morning before CBD/vehicle administration (i.e. assessment occurred roughly 24 hours after the last drug administration and CBD should have been mostly metabolized by then). Our results are in line with findings from Uttl et al. (2021), who similarly found that a single dose of 10 or 60 mg/kg CBD i.p. administered to PD12 rats did not influence initial performance on the bar holding test but impaired performance improvement 24 hours later, potentially a sign of impaired motor learning. Of note, the present study found no effect of CBD on negative geotaxis latency which was assessed in the days following righting reflex testing, which in turn indicates that CBD’s effects in this regard might be temporary and there are no lasting effects on neuromotor performance. In line with this, there was also no effect of CBD (or pain) on body weight gain until weaning or age at eye opening or sexual maturity, suggesting that neither CBD nor pain were strong enough stimuli to interfere with these essential milestones.

### Neither Pain nor CBD Affected Adult Pain Sensitivity or Object

#### Recognition Memory

There were no significant effects from neonatal pain and/or CBD on thermal pain threshold, as measured by paw-lick latency in the hot plate test. Many previous studies have also reported no significant effect of neonatal pain on adult pain sensitivity (Anand et al., 1999; Davis et al., 2021; Page et al., 2013; Ranger et al., 2018; van den Hoogen et al., 2020) but some studies have reported that neonatal procedural pain exposure increases adult sensitivity to pain (Johnston & Walker, 2003; Min et al., 2022; Nuseir et al., 2017; Page et al., 2013; Sanada et al., 2014; van den Hoogen et al., 2021; van den Hoogen et al., 2020). It is likely that experimental details such as timing of exposure, intensity and type of the painful stimulus or even experimenter gender mediate the effects of neonatal pain on adult pain sensitivity (Sorge et al., 2014).

The present study also found no effect of neonatal CBD and/or procedural pain exposure on adult recognition memory in the novel object preference test. However, it is possible that the stimuli we used for this test were too similar, as it appears that control touch + vehicle subjects did not consistently recognize when the stimuli changed. In other words, our test may have been too difficult even for “healthy” rats, and thus our results cannot be used to draw conclusions about the impact of CBD or neonatal pain on performances in this test. Previous studies have found that higher doses of CBD (50 mg/kg i.p. given on PND 1,3 and 5) can impact adult spatial memory in adult female rats in the Barnes maze, while males did not display any deficits (Wadhwa et al., 2024), suggesting that more research is needed to better understand the possible sexually-dimorphic impact of neonatal CBD exposure on later cognitive function.

### Neonatal Pain Increases Adult Anxiety-like Behavior

Neonatal pain exposure resulted in males spending less time in the open arms of the EZM as well as spending less time and making fewer entries into the center of the open field test, suggesting that neonatal pain exposure increases anxiety-like behavior in male rats. Interestingly, a similar trend/pattern was seen in female rats, but only in the 10 mg/kg CBD dose group with pain females receiving 10 mg/kg CBD spending less time and making fewer entries into the open arms of the EZM compared to touch females receiving 10mg/kg, while an opposite pattern was seen for the 5 mg/kg females (with Pain females making slightly, but non-significantly more entries compared to touch females (p=0.062)). A similar drug by pain interaction was also seen in males for the overall distance travelled with 10 mg/kg pain males travelling less than the 10 mg/kg touch males. This indicates that there is a complex sex- and dose-dependent effect on how neonatal pain and CBD exposure influences adult anxiety-like and exploratory behavior.

This increased anxiety-like behavior is in line with previous studies demonstrating that “mild” neonatal pain exposure can influence later-life anxiety-like behavior (Page et al., 2005; Schellinck et al., 2003). Interestingly, exposure to more frequent or more severe neonatal pain does not seem to influence adult anxiety-like behavior (de Kort et al., 2021, 2022; Mooney-Leber & Brummelte, 2020; Page et al., 2005; Ranger et al., 2018; Zuke et al., 2019). This was best demonstrated by Page et al. (2005) who reported that 4 needle pokes per day every other day from PD0-7 (i.e., a relatively mild stressor and similar to our pain exposure used here) increased adult anxiety-like behavior in the elevated plus maze, but 4 pokes/day administered every day for 8 days did not change adult behavior in the EPM. Similarly, our previous study using repetitive needle pokes that dissipated from 8 pokes on postnatal day 1 to 4 pokes on postnatal day 4 (i.e. more pokes than in the current study), did not find a pain effect for the time in the center of the open field. This suggests that neonatal pain exposure has a very unique “dose-dependent” impact on adult anxiety-like behavior that deserves further investigation. This is particularly important because human studies suggest that former-preterm children display increased internalizing behaviors and increased risk for anxiety disorders which is associated with more procedural pain exposure in the NICU (Ranger & Grunau, 2014; Victoria & Murphy, 2016).

Pretreatment with 5 or 10 mg/kg CBD was unable to prevent changes in adult anxiety-like behavior in the present study though some interaction effects suggest that CBD treatment may modify, albeit not rescue, the effects of pain. Rego et al. (2024) similarly found intraperitoneal pretreatment with 5 or 20 mg/kg CBD did not prevent neonatal inflammatory pain from increasing adult anxiety-like behavior. Even 50 mg/kg CBD given neonatally did not alter behavior in the elevated plus maze in adult rats (Wadhwa et al., 2024-09-10), though this dose resulted in spatial memory impairments and changes to dendritic morphology in young adulthood. This suggests that neonatal CBD treatment may not be an efficient pain management strategy and emphasizes the need for future research on alternative methods of pain management that can mitigate the long-term consequences of acute pain exposure on adult anxiety-like behavior and other adverse consequences.

### Neonatal CBD Increases Baseline CORT in Males and Decreased Body Weight in Females at Adulthood

Adult male rats exposed to 10 mg/kg CBD neonatally had slightly higher baseline CORT levels than vehicle male rats. This effect is in line with previous research showing that neonatal exposure to a CB1R agonist is also associated with increased baseline CORT in adolescent rats (Llorente et al., 2007). Further, adolescent THC exposure is linked to increased baseline CORT in adult rats (Ferland et al., 2023), suggesting that early-life manipulations of the ECS can result in long-lasting changes to the HPA axis and stress sensitivity.

On the other hand, neonatal procedural pain 4 times a day from PD2-5 did not significantly alter adult CORT responses to restraint stress in the current study. This is in slight contrast to our previous study, where we found that pain-exposed adult females (8 pokes on PD1, 6 pokes on PD2, and 4 pokes on PD3-4) had higher CORT recovery levels (i.e. CORT was still elevated 60 minutes after the stressor) compared to touch female rats (Mooney-Leber & Brummelte, 2020). It is noteworthy that pups in our previous study received almost twice as many painful paw pricks, and this may have been a more powerful stressor to induce long-term changes in HPA axis function, which is in line with our findings for pain inducing increased anxiety-like behavior in a dose-dependent manner (as discussed above). Additionally, it is also important to note that all pups in the current study received repeated neonatal gavage from PD 2-5, which may have had its own impact on stress regulation and could thus explain the slight discrepancy to our previous findings.

Interestingly, we found that neonatal CBD decreased adult body weight in female subjects but had no effect on developmental weight gain. In contrast, gestational or postnatal exposure to other cannabinoids or CB receptor agonists can alter early-life weight gain (Antonelli et al., 2005; Chen et al., 2016; Dalterio et al., 1984; Iezzi et al., 2022; Llorente et al., 2007; Philippot et al., 2016). The alteration of female adult body weight in the present study without changes in developmental body weight or male adult body weight suggest that the effect of neonatal CBD exposure was sex-specific and may involve long-term alterations in metabolism or body-weight regulation. Additionally, our results suggest the effect of neonatal CBD on adult female body weight might be modulated by neonatal pain exposure, however, the follow-up analysis of the interaction effect did not reveal a clear direction for this relationship. Further research is necessary to determine the mechanisms of action behind both effects.

## Conclusion

The present study examined if CBD could be an effective form of pain management for neonates experiencing neonatal pain-related stress using a translational rat model. Although CBD successfully decreased USVs in neonatal pups, pain also decreased USVs, and the observed reduction of USVs from both experimental manipulations makes interpretation slightly difficult. Reduced USVs may indicate reduced stress levels, as anxiolytic drugs can decrease these vocalizations (Winslow & Insel, 1991), however, the fact that pups also vocalized less after pain exposure indicates there is a complex relationship between stress states and USVs. CBD also transiently impaired neuromotor reflex improvement, increased baseline CORT levels in adult males and lowered body weight in adult females. On the other hand, this study found no significant effect of neonatal exposure to CBD (and/or repeated procedural pain) on neonatal body weight, age at sexual maturity in male and female rats, or estrus cycling of adult female subjects. Previous studies suggested that CBD is a reproductive toxin for adult male rodents, and can acutely reduce testosterone production, decrease testis size, prevent spermatogenesis, and reduce fertility (as reviewed by Carvalho et al., 2020). However, there is limited research on whether neonatal exposure to CBD also alters later-life reproductive function. The lack of research on how later-life reproductive function is influenced by early-life CBD exposure is especially problematic because CBD (as the main ingredient of Epidiolex) is currently FDA-approved and used to treat intractable forms of epilepsy in children 2 years of age and older. Previous studies have demonstrated that the consequences of an individual neonatal or gestational stressor can be heavily influenced by other perinatal stressors (Davis & Burman, 2021; Mooney-Leber, 2018; Mooney-Leber & Brummelte, 2020; Timmerman et al., 2021) and we cannot rule out the possibility that the stress of oral gavage could have altered the effects of neonatal procedural pain. However, oral administration is translationally relevant because Epidiolex, the CBD-containing drug used in the present study, is also administered orally to human patients. Further, preterm infants have to go through a multitude of stressors while in the NICU, and a treatment for neonatal pain that becomes ineffective in the presence of other stressors is therefore unlikely to be of practical use.

Given that over 10% of all live births in the US are preterm, and preterm infants display a multitude of neurodevelopmental changes associated with the painful procedures they experience in the NICU, it is imperative to use effective pain management strategies that are able to prevent the short- and long-term effects of these necessary painful exposures. More research is needed to further investigate whether CBD may be effective for early-life pain management, maybe in combination with other medications (Rego et al., 2024) or whether it may have detrimental effects on neurodevelopment or reproductive function later in life.

## Acknowledgements

This work was supported by a Betty Neitzel Faculty Award to SB. We would like to thank Kit Tran, Armina Khan, LaShawna Muhammad, Lauren Richardson, Abigail Myers, Jecenia Duran, Hanna Cox, Rubina Khan, Auttum Haseltine, Adela Veladzic, for their help with this work.

## Reference list

Adams, J., & Jones, S. M. (1984). Age differences in water maze performance and swimming behavior in the rat. Physiol Behav, 33(6), 851–855. 10.1016/0031-9384(84)90218-x

Akingbasote, J., Szlapinski, S., Charrette, A., Hilmas, C. J., & Guthrie, N. (2022). Safety of cannabis- and hemp-derived constituents in reproduction and development. In Reproductive and Developmental Toxicology (pp. 455–487). 10.1016/b978-0-323-89773-0.00024-2

Anand, K. J., Coskun, V., Thrivikraman, K. V., Nemeroff, C. B., & Plotsky, P. M. (1999). Long-term behavioral effects of repetitive pain in neonatal rat pups. Physiol Behav, 66(4), 627–637. 10.1016/s0031-9384(98)00338-2

Anand, K. J., Garg, S., Rovnaghi, C. R., Narsinghani, U., Bhutta, A. T., & Hall, R. W. (2007). Ketamine reduces the cell death following inflammatory pain in newborn rat brain. Pediatr Res, 62(3), 283–290. 10.1203/PDR.0b013e3180986d2f

Antonelli, T., Tomasini, M. C., Tattoli, M., Cassano, T., Tanganelli, S., Finetti, S.,…Ferraro, L. (2005). Prenatal exposure to the CB1 receptor agonist WIN 55,212-2 causes learning disruption associated with impaired cortical NMDA receptor function and emotional reactivity changes in rat offspring. Cereb Cortex, 15(12), 2013–2020. 10.1093/cercor/bhi076

Benoit, J. D., Rakic, P., & Frick, K. M. (2015). Prenatal stress induces spatial memory deficits and epigenetic changes in the hippocampus indicative of heterochromatin formation and reduced gene expression. Behav Brain Res, 281, 1–8. 10.1016/j.bbr.2014.12.001

Blazevic, S., Merkler, M., Persic, D., & Hranilovic, D. (2017). Chronic postnatal monoamine oxidase inhibition affects affiliative behavior in rat pups. Pharmacol Biochem Behav, 153, 60–68. 10.1016/j.pbb.2016.12.008

Boulanger, J. J., & Messier, C. (2017). Unbiased stereological analysis of the fate of oligodendrocyte progenitor cells in the adult mouse brain and effect of reference memory training. Behav Brain Res, 329, 127–139. 10.1016/j.bbr.2017.04.027

Boulanger-Bertolus, J., Rincon-Cortes, M., Sullivan, R. M., & Mouly, A. M. (2017). Understanding pup affective state through ethologically significant ultrasonic vocalization frequency. Sci Rep, 7(1), 13483. 10.1038/s41598-017-13518-6

Branchi, I., Santucci, D., & Alleva, E. (2001). Ultrasonic vocalisation emitted by infant rodents: a tool for assessment of neurobehavioural development. Behavioural Brain Research, 125(1-2), 49–56. 10.1016/s0166-4328(01)00277-7

Brummelte, S., Chau, C. M., Cepeda, I. L., Degenhardt, A., Weinberg, J., Synnes, A. R., & Grunau, R. E. (2015). Cortisol levels in former preterm children at school age are predicted by neonatal procedural pain-related stress. Psychoneuroendocrinology, 51, 151–163. 10.1016/j.psyneuen.2014.09.018

Brummelte, S., Grunau, R. E., Chau, V., Poskitt, K. J., Brant, R., Vinall, J.,…Miller, S. P. (2012). Procedural pain and brain development in premature newborns. Ann Neurol, 71(3), 385–396. 10.1002/ana.22267

Butkevich, I. P., Mikhailenko, V. A., & Vershinina, E. A. (2023). Long-Term Influences of Neonatal Pain-Related Stress on Cognitive and Stress-Hormonal Functions in Rats: Age and Sex Aspects. Journal of Evolutionary Biochemistry and Physiology, 59(3), 756–768. 10.1134/s0022093023030109

Campbell-Yeo, M. L., Disher, T. C., Benoit, B. L., & Johnston, C. C. (2015). Understanding kangaroo care and its benefits to preterm infants. Pediatric Health Med Ther, 6, 15–32. 10.2147/PHMT.S51869

Carbajal, R., Rousset, A., Danan, C., Coquery, S., Nolent, P., Ducrocq, S.,…Breart, G. (2008). Epidemiology and treatment of painful procedures in neonates in intensive care units. JAMA, 300(1), 60–70. 10.1001/jama.300.1.60

Carson, R., Monaghan-Nichols, A. P., DeFranco, D. B., & Rudine, A. C. (2016). Effects of antenatal glucocorticoids on the developing brain. Steroids, 114, 25–32. 10.1016/j.steroids.2016.05.012

Carvalho, R. K., Andersen, M. L., & Mazaro-Costa, R. (2020). The effects of cannabidiol on male reproductive system: A literature review. J Appl Toxicol, 40(1), 132–150. 10.1002/jat.3831

Catalani, A., Marinelli, M., Scaccianoce, S., Nicolai, R., Muscolo, L. A. A., Porcu, A.,…Angelucci, L. (1993). Progeny of mothers drinking corticosterone during lactation has lower stress-induced corticosterone secretion and better cognitive performance. Brain Research, 624(1-2), 209–215. 10.1016/0006-8993(93)90079-3

Chen, M., Xia, D., Min, C., Zhao, X., Chen, Y., Liu, L., & Li, X. (2016). Neonatal repetitive pain in rats leads to impaired spatial learning and dysregulated hypothalamic-pituitary-adrenal axis function in later life. Sci Rep, 6, 39159. 10.1038/srep39159

Coffey, E. B. J., Nicol, T., White-Schwoch, T., Chandrasekaran, B., Krizman, J., Skoe, E.,…Kraus, N. (2019-11-06). Evolving perspectives on the sources of the frequency-following response. Nature Communications 2019 10:1, 10(1). 10.1038/s41467-019-13003-w

Corcoran, L., Roche, M., & Finn, D. P. (2015). The Role of the Brain’s Endocannabinoid System in Pain and Its Modulation by Stress. Int Rev Neurobiol, 125, 203–255. 10.1016/bs.irn.2015.10.003

Corsi, D. J., Murphy, M. S. Q., & Cook, J. (2021). The Effects of Cannabis on Female Reproductive Health Across the Life Course. Cannabis Cannabinoid Res, 6(4), 275–287. 10.1089/can.2020.0065

Costa, B., Trovato, A. E., Comelli, F., Giagnoni, G., & Colleoni, M. (2007). The non-psychoactive cannabis constituent cannabidiol is an orally effective therapeutic agent in rat chronic inflammatory and neuropathic pain. Eur J Pharmacol, 556(1-3), 75–83. 10.1016/j.ejphar.2006.11.006

Dalterio, S., Steger, R., Mayfield, D., & Bartke, A. (1984). Early cannabinoid exposure influences neuroendocrine and reproductive functions in male mice: I. Prenatal exposure. Pharmacol Biochem Behav, 20(1), 107–113. 10.1016/0091-3057(84)90110-2

Davis, S. M., & Burman, M. A. (2021). Maternal separation with neonatal pain influences later-life fear conditioning and somatosenation in male and female rats. Stress, 24(5), 504–513. 10.1080/10253890.2020.1825674

Davis, S. M., Zuke, J. T., Berchulski, M. R., & Burman, M. A. (2021). Amygdalar Corticotropin-Releasing Factor Signaling Is Required for Later-Life Behavioral Dysfunction Following Neonatal Pain. Front Physiol, 12, 660792. 10.3389/fphys.2021.660792

de Kort, A. R., Joosten, E. A., Patijn, J., Tibboel, D., & van den Hoogen, N. J. (2021). Neonatal procedural pain affects state, but not trait anxiety behavior in adult rats. Dev Psychobiol, 63(8), e22210. 10.1002/dev.22210

de Kort, A. R., Joosten, E. A., Patijn, J., Tibboel, D., & van den Hoogen, N. J. (2022). Selective Targeting of Serotonin 5-HT1a and 5-HT3 Receptors Attenuates Acute and Long-Term Hypersensitivity Associated With Neonatal Procedural Pain. Front Pain Res (Lausanne), 3, 872587. 10.3389/fpain.2022.872587

De Vita, M. J., Maisto, S. A., Gilmour, C. E., McGuire, L., Tarvin, E., & Moskal, D. (2021). The effects of cannabidiol and analgesic expectancies on experimental pain reactivity in healthy adults: A balanced placebo design trial. Experimental and Clinical Psychopharmacology. 10.1037/pha0000465

De Vita, M. J., Moskal, D., Maisto, S. A., & Ansell, E. B. (2018). Association of Cannabinoid Administration With Experimental Pain in Healthy Adults: A Systematic Review and Meta-analysis. JAMA Psychiatry, 75(11), 1118–1127. 10.1001/jamapsychiatry.2018.2503

Doesburg, S. M., Moiseev, A., Herdman, A. T., Ribary, U., & Grunau, R. E. (2013/11/15). Frontiers | Region-Specific Slowing of Alpha Oscillations is Associated with Visual-Perceptual Abilities in Children Born Very Preterm. Frontiers in Human Neuroscience, 7. 10.3389/fnhum.2013.00791

Feldman, R., Rosenthal, Z., & Eidelman, A. I. (2014). Maternal-preterm skin-to-skin contact enhances child physiologic organization and cognitive control across the first 10 years of life. Biol Psychiatry, 75(1), 56–64. 10.1016/j.biopsych.2013.08.012

Ferland, J. N., Ellis, R. J., Rompala, G., Landry, J. A., Callens, J. E., Ly, A.,…Hurd, Y. L. (2023). Dose mediates the protracted effects of adolescent THC exposure on reward and stress reactivity in males relevant to perturbation of the basolateral amygdala transcriptome. Mol Psychiatry, 28(6), 2583–2593. 10.1038/s41380-022-01467-0

Fonseca, A., Sanata, G., Bosque Ortiz, G., Bampi, S., Dietrich, M., (2021). Analysis of ultrasonic vocalizations from mice using computer vision and machine learning. eLife, 10. 10.7554/eLife.59161

Fride, E. (2008). Multiple roles for the endocannabinoid system during the earliest stages of life: pre- and postnatal development. J Neuroendocrinol, 20 Suppl 1, 75–81. 10.1111/j.1365-2826.2008.01670.x

Fukumoto, K., Morita, T., Mayanagi, T., Tanokashira, D., Yoshida, T., Sakai, A., & Sobue, K. (2009). Detrimental effects of glucocorticoids on neuronal migration during brain development. Mol Psychiatry, 14(12), 1119–1131. 10.1038/mp.2009.60

Gazendam, A., Nucci, N., Gouveia, K., Abdel Khalik, H., Rubinger, L., & Johal, H. (2020). Cannabinoids in the Management of Acute Pain: A Systematic Review and Meta-analysis. Cannabis Cannabinoid Res, 5(4), 290–297. 10.1089/can.2019.0079

Grant, K. S., Petroff, R., Isoherranen, N., Stella, N., & Burbacher, T. M. (2018). Cannabis use during pregnancy: Pharmacokinetics and effects on child development. Pharmacol Ther, 182, 133–151. 10.1016/j.pharmthera.2017.08.014

Grimsley, J. M. S., Monaghan, J. J. M., & Wenstrup, J. J. (Mar 9, 2011). Development of Social Vocalizations in Mice. PLOS ONE, 6(3). 10.1371/journal.pone.0017460

Grunau, R. E. (2013). Neonatal pain in very preterm infants: long-term effects on brain, neurodevelopment and pain reactivity. Rambam Maimonides Med J, 4(4), e0025. 10.5041/RMMJ.10132

Grunau, R. E., Cepeda, I. L., Chau, C. M., Brummelte, S., Weinberg, J., Lavoie, P. M.,…Turvey, S. E. (2013). Neonatal pain-related stress and NFKBIA genotype are associated with altered cortisol levels in preterm boys at school age. PLoS One, 8(9), e73926. 10.1371/journal.pone.0073926

Grunau, R. E., Holsti, L., Haley, D. W., Oberlander, T., Weinberg, J., Solimano, A.,…Yu, W. (2005). Neonatal procedural pain exposure predicts lower cortisol and behavioral reactivity in preterm infants in the NICU. Pain, 113(3), 293–300. 10.1016/j.pain.2004.10.020

Grunau, R. E., Whitfield, M. F., Petrie-Thomas, J., Synnes, A. R., Cepeda, I. L., Keidar, A.,…Johannesen, D. (2009). Neonatal pain, parenting stress and interaction, in relation to cognitive and motor development at 8 and 18 months in preterm infants. Pain, 143(1-2), 138–146. 10.1016/j.pain.2009.02.014

Naito, H., & Tonoue, T., (1987). Sex difference in ultrasound distress call by rat pups. Behavioural Brain Research, 25(1). 10.1016/0166-4328(87)90041-6

Hammell, D. C., Zhang, L. P., Ma, F., Abshire, S. M., McIlwrath, S. L., Stinchcomb, A. L., & Westlund, K. N. (2016). Transdermal cannabidiol reduces inflammation and pain-related behaviours in a rat model of arthritis. Eur J Pain, 20(6), 936–948. 10.1002/ejp.818

Harkany, T., Keimpema, E., Barabas, K., & Mulder, J. (2008). Endocannabinoid functions controlling neuronal specification during brain development. Mol Cell Endocrinol, 286(1-2 Suppl 1), S84-90. 10.1016/j.mce.2008.02.011

Hill, M. N., & Tasker, J. G. (2012). Endocannabinoid signaling, glucocorticoid-mediated negative feedback, and regulation of the hypothalamic-pituitary-adrenal axis. Neuroscience, 204, 5–16. 10.1016/j.neuroscience.2011.12.030

Iezzi, D., Caceres-Rodriguez, A., Chavis, P., & Manzoni, O. J. J. (2022). In utero exposure to cannabidiol disrupts select early-life behaviors in a sex-specific manner. Transl Psychiatry, 12(1), 501. 10.1038/s41398-022-02271-8

Ise, S., & Ohta, H. (2009). Power spectrum analysis of ultrasonic vocalization elicited by maternal separation in rat pups. Brain Res, 1283, 58–64. 10.1016/j.brainres.2009.06.003

Johnston, C. C., Filion, F., Campbell-Yeo, M., Goulet, C., Bell, L., McNaughton, K.,…Walker, C. D. (2008). Kangaroo mother care diminishes pain from heel lance in very preterm neonates: a crossover trial. BMC Pediatr, 8, 13. 10.1186/1471-2431-8-13

Johnston, C. C., & Walker, C. D. (2003). The effects of exposure to repeated minor pain during the neonatal period on formalin pain behaviour and thermal withdrawal latencies. Pain Res Manag, 8(4), 213–217. 10.1155/2003/305409

Kwok, C. H., Devonshire, I. M., Imraish, A., Greenspon, C. M., Lockwood, S., Fielden, C.,…Hathway, G. J. (2017). Age-dependent plasticity in endocannabinoid modulation of pain processing through postnatal development. Pain, 158(11), 2222–2232. 10.1097/j.pain.0000000000001027

Lehmann, J., Pryce, C. R., Jongen-Relo, A. L., Stohr, T., Pothuizen, H. H., & Feldon, J. (2002). Comparison of maternal separation and early handling in terms of their neurobehavioral effects in aged rats. Neurobiol Aging, 23(3), 457–466. 10.1016/s0197-4580(01)00320-7

Llorente, R., Arranz, L., Marco, E. M., Moreno, E., Puerto, M., Guaza, C.,…Viveros, M. P. (2007). Early maternal deprivation and neonatal single administration with a cannabinoid agonist induce long-term sex-dependent psychoimmunoendocrine effects in adolescent rats. Psychoneuroendocrinology, 32(6), 636–650. 10.1016/j.psyneuen.2007.04.002

Malheiros, J. M., Lima, M., Avanzi, R. D., Gomes da Silva, S., Suchecki, D., Guinsburg, R., & Covolan, L. (2014). Repetitive noxious neonatal stimuli increases dentate gyrus cell proliferation and hippocampal brain-derived neurotrophic factor levels. Hippocampus, 24(4), 415–423. 10.1002/hipo.22235

Manduca, A., Campolongo, P., & Trezza, V. (2012). Cannabinoid modulation of mother-infant interaction: is it just about milk? Rev Neurosci, 23(5-6), 707–722. 10.1515/revneuro-2012-0074

Manduca, A., Servadio, M., Melancia, F., Schiavi, S., Manzoni, O. J., & Trezza, V. (2020). Sex-specific behavioural deficits induced at early life by prenatal exposure to the cannabinoid receptor agonist WIN55, 212-2 depend on mGlu5 receptor signalling. Br J Pharmacol, 177(2), 449–463. 10.1111/bph.14879

McCormick, C. M., Rioux, T., Fisher, R., Lang, K., MacLaury, K., & Teillon, S. M. (2001). Effects of neonatal corticosterone treatment on maze performance and HPA axis in juvenile rats. Physiol Behav, 74(3), 371–379. 10.1016/s0031-9384(01)00574-1

McLean, A. C., Valenzuela, N., Fai, S., & Bennett, S. A. (2012). Performing vaginal lavage, crystal violet staining, and vaginal cytological evaluation for mouse estrous cycle staging identification. J Vis Exp(67), e4389. 10.3791/4389

McPherson, C., & Grunau, R. E. (2014). Neonatal pain control and neurologic effects of anesthetics and sedatives in preterm infants. Clin Perinatol, 41(1), 209–227. 10.1016/j.clp.2013.10.002

Min, C., Ling, R., Chen, M., Xia, D., Chen, R., & Li, X. (2022). Enriched environment rescues neonatal pain induced cognitive deficits and the impaired hippocampal synaptic plasticity later in life. Dev Neurobiol, 82(6), 545–561. 10.1002/dneu.22898

Mlost, J., Bryk, M., & Starowicz, K. (2020). Cannabidiol for Pain Treatment: Focus on Pharmacology and Mechanism of Action. Int J Mol Sci, 21(22). 10.3390/ijms21228870

Modir, F., Elahdadi Salmani, M., Goudarzi, I., Lashkarboluki, T., & Abrari, K. (2014). Prenatal stress decreases spatial learning and memory retrieval of the adult male offspring of rats. Physiol Behav, 129, 104–109. 10.1016/j.physbeh.2014.02.040

Mooney-Leber, S. M. (2018). The Impact Of Neonatal Pain And Reduced Maternal Care On Brain And Behavioral Development. Wayne State University Dissertations, 1948.

Mooney-Leber, S. M., & Brummelte, S. (2020). Neonatal pain and reduced maternal care alter adult behavior and hypothalamic-pituitary-adrenal axis reactivity in a sex-specific manner. Dev Psychobiol, 62(5), 631–643. 10.1002/dev.21941

Mooney-Leber, S. M., Spielmann, S. S., & Brummelte, S. (2018). Repetitive neonatal pain and reduced maternal care alter brain neurochemistry. Dev Psychobiol, 60(8), 963–974. 10.1002/dev.21777

Moore, E. R., Anderson, G. C., Bergman, N., & Dowswell, T. (2012). Early skin-to-skin contact for mothers and their healthy newborn infants. Cochrane Database Syst Rev(5), CD003519. 10.1002/14651858.CD003519.pub3

Morelius, E., Ortenstrand, A., Theodorsson, E., & Frostell, A. (2015). A randomised trial of continuous skin-to-skin contact after preterm birth and the effects on salivary cortisol, parental stress, depression, and breastfeeding. Early Hum Dev, 91(1), 63–70. 10.1016/j.earlhumdev.2014.12.005

Nuseir, K. Q., Alzoubi, K. H., Alhusban, A., Bawaane, A., Al-Azzani, M., & Khabour, O. F. (2017). Sucrose and naltrexone prevent increased pain sensitivity and impaired long-term memory induced by repetitive neonatal noxious stimulation: Role of BDNF and beta-endorphin. Physiol Behav, 179, 213–219. 10.1016/j.physbeh.2017.06.015

Ohlsson, A., & Shah, P. S. (2020). Paracetamol (acetaminophen) for prevention or treatment of pain in newborns. Cochrane Database Syst Rev, 1, CD011219. 10.1002/14651858.CD011219.pub4

Oller, D. K., Griebel, U., Bowman, D. D., Bene, E., Long, H. L., Yoo, H., & Ramsay, G. (2020). Infant boys are more vocal than infant girls. Curr Biol, 30(10), R426–R427. 10.1016/j.cub.2020.03.049

Page, G. G., Blakely, W. P., & Kim, M. (2005). The impact of early repeated pain experiences on stress responsiveness and emotionality at maturity in rats. Brain Behav Immun, 19(1), 78–87. 10.1016/j.bbi.2004.05.002

Page, G. G., Hayat, M. J., & Kozachik, S. L. (2013). Sex differences in pain responses at maturity following neonatal repeated minor pain exposure in rats. Biol Res Nurs, 15(1), 96–104. 10.1177/1099800411419493

Papadimitriou, A., & Priftis, K. N. (2009). Regulation of the hypothalamic-pituitary-adrenal axis. Neuroimmunomodulation, 16(5), 265–271. 10.1159/000216184

Philippot, G., Nyberg, F., Gordh, T., Fredriksson, A., & Viberg, H. (2016). Short-term exposure and long-term consequences of neonatal exposure to Delta(9)-tetrahydrocannabinol (THC) and ibuprofen in mice. Behav Brain Res, 307, 137–144. 10.1016/j.bbr.2016.04.001

Plotsky, P. M., Thrivikraman, K. V., Nemeroff, C. B., Caldji, C., Sharma, S., & Meaney, M. J. (2005). Long-term consequences of neonatal rearing on central corticotropin-releasing factor systems in adult male rat offspring. Neuropsychopharmacology, 30(12), 2192–2204. 10.1038/sj.npp.1300769

Ranger, M., & Grunau, R. E. (2014). Early repetitive pain in preterm infants in relation to the developing brain. Pain Manag, 4(1), 57–67. 10.2217/pmt.13.61

Ranger, M., Synnes, A. R., Vinall, J., & Grunau, R. E. (2014). Internalizing behaviours in school-age children born very preterm are predicted by neonatal pain and morphine exposure. Eur J Pain, 18(6), 844–852. 10.1002/j.1532-2149.2013.00431.x

Ranger, M., Tremblay, S., Chau, C. M. Y., Holsti, L., Grunau, R. E., & Goldowitz, D. (2018). Adverse Behavioral Changes in Adult Mice Following Neonatal Repeated Exposure to Pain and Sucrose. Front Psychol, 9, 2394. 10.3389/fpsyg.2018.02394

Rego, D. S. B., Calio, M. L., Filev, R., Mello, L. E., & Leslie, A. (2024). Long-term Effects of Cannabidiol and/or Fentanyl Exposure in Rats Submitted to Neonatal Pain. J Pain, 25(3), 715–729. 10.1016/j.jpain.2023.10.001

Rubino, T., & Parolaro, D. (2008). Long lasting consequences of cannabis exposure in adolescence. Mol Cell Endocrinol, 286(1-2 Suppl 1), S108–113. 10.1016/j.mce.2008.02.003

Sanada, L. S., Sato, K. L., Machado, N. L., Carmo Ede, C., Sluka, K. A., & Fazan, V. P. (2014). Cortex glial cells activation, associated with lowered mechanical thresholds and motor dysfunction, persists into adulthood after neonatal pain. Int J Dev Neurosci, 35, 55–63. 10.1016/j.ijdevneu.2014.03.008

Schellinck, H. M., Stanford, L., & Darrah, M. (2003). Repetitive acute pain in infancy increases anxiety but does not alter spatial learning ability in juvenile mice. Behavioural Brain Research, 142(1-2), 157–165. 10.1016/s0166-4328(02)00406-0

Scheyer, A. F., Borsoi, M., Wager-Miller, J., Pelissier-Alicot, A. L., Murphy, M. N., Mackie, K., & Manzoni, O. J. J. (2020). Cannabinoid Exposure via Lactation in Rats Disrupts Perinatal Programming of the Gamma-Aminobutyric Acid Trajectory and Select Early-Life Behaviors. Biol Psychiatry, 87(7), 666–677. 10.1016/j.biopsych.2019.08.023

Schneider, M. (2009). Cannabis use in pregnancy and early life and its consequences: animal models. Eur Arch Psychiatry Clin Neurosci, 259(7), 383–393. 10.1007/s00406-009-0026-0

Schwaller, F., & Fitzgerald, M. (2014). The consequences of pain in early life: injury-induced plasticity in developing pain pathways. Eur J Neurosci, 39(3), 344–352. 10.1111/ejn.12414

Simola, N. (2015). Rat Ultrasonic Vocalizations and Behavioral Neuropharmacology: From the Screening of Drugs to the Study of Disease. Curr Neuropharmacol, 13(2), 164–179. 10.2174/1570159x13999150318113800

Slater, R., Cornelissen, L., Fabrizi, L., Patten, D., Yoxen, J., Worley, A.,…Fitzgerald, M. (2010). Oral sucrose as an analgesic drug for procedural pain in newborn infants: a randomised controlled trial. Lancet, 376(9748), 1225–1232. 10.1016/S0140-6736(10)61303-7

Sorge, R. E., Martin, L. J., Isbester, K. A., Sotocinal, S. G., Rosen, S., Tuttle, A. H.,…Mogil, J. S. (2014). Olfactory exposure to males, including men, causes stress and related analgesia in rodents. Nat Methods, 11(6), 629–632. 10.1038/nmeth.2935

Takahashi, L. K. (1992). Developmental expression of defensive responses during exposure to conspecific adults in preweanling rats (Rattus norvegicus). J Comp Psychol, 106(1), 69–77. 10.1037/0735-7036.106.1.69

Takahashi, L. K., Turner, J. G., & Kalin, N. H. (1991). Development of stress-induced responses in preweanling rats. Dev Psychobiol, 24(5), 341–360. 10.1002/dev.420240504

Timmerman, B. M., Mooney-Leber, S. M., & Brummelte, S. (2021). The effects of neonatal procedural pain and maternal isolation on hippocampal cell proliferation and reelin concentration in neonatal and adult male and female rats. Dev Psychobiol, 63(8), e22212. 10.1002/dev.22212

Trezza, V., Campolongo, P., Manduca, A., Morena, M., Palmery, M., Vanderschuren, L. J., & Cuomo, V. (2012). Altering endocannabinoid neurotransmission at critical developmental ages: impact on rodent emotionality and cognitive performance. Front Behav Neurosci, 6, 2. 10.3389/fnbeh.2012.00002

Uttl, L., Hlozek, T., Mares, P., Palenicek, T., & Kubova, H. (2021). Anticonvulsive Effects and Pharmacokinetic Profile of Cannabidiol (CBD) in the Pentylenetetrazol (PTZ) or N-Methyl-D-Aspartate (NMDA) Models of Seizures in Infantile Rats. Int J Mol Sci, 23(1). 10.3390/ijms23010094

Valeri, B. O., Ranger, M., Chau, C. M., Cepeda, I. L., Synnes, A., Linhares, M. B., & Grunau, R. E. (2016). Neonatal Invasive Procedures Predict Pain Intensity at School Age in Children Born Very Preterm. Clin J Pain, 32(12), 1086–1093. 10.1097/AJP.0000000000000353

van den Hoogen, N. J., de Geus, T. J., Patijn, J., Tibboel, D., & Joosten, E. A. (2021). Methadone effectively attenuates acute and long-term consequences of neonatal repetitive procedural pain in a rat model. Pediatr Res. 10.1038/s41390-020-01353-x

van den Hoogen, N. J., Patijn, J., Tibboel, D., & Joosten, E. A. (2020). Repetitive noxious stimuli during early development affect acute and long-term mechanical sensitivity in rats. Pediatr Res, 87(1), 26–31. 10.1038/s41390-019-0420-x

Victoria, N. C., & Murphy, A. Z. (2016). Exposure to Early Life Pain: Long Term Consequences and Contributing Mechanisms. Curr Opin Behav Sci, 7, 61–68. 10.1016/j.cobeha.2015.11.015

Wadhwa, M., Chinn, G. A., Sasaki Russell, J. M., Hellman, J., & Sall, J. W. (2024-09-10). Neonatal Cannabidiol Exposure Impairs Spatial Memory and Disrupts Neuronal Dendritic Morphology in Young Adult Rats. Cannabis and Cannabinoid Research. 10.1089/can.2024.0010

Walker, S. M. (2014). Neonatal pain. Paediatr Anaesth, 24(1), 39–48. 10.1111/pan.12293

Wanner, N. M., Colwell, M., Drown, C., & Faulk, C. (2021). Developmental cannabidiol exposure increases anxiety and modifies genome-wide brain DNA methylation in adult female mice. Clin Epigenetics, 13(1), 4. 10.1186/s13148-020-00993-4

Wiedenmayer, C. P., Lyo, D., & Barr, G. A. (2003). Rat pups reduce ultrasonic vocalization after exposure to an adult male rat. Dev Psychobiol, 42(4), 386–391. 10.1002/dev.10112

Williams, M. D., & Lascelles, B. D. X. (2020). Early Neonatal Pain-A Review of Clinical and Experimental Implications on Painful Conditions Later in Life. Front Pediatr, 8, 30. 10.3389/fped.2020.00030

Winslow, J. T., & Insel, T. R. (1991/01/01). Infant rat separation is a sensitive test for novel anxiolytics. Progress in Neuro-Psychopharmacology and Biological Psychiatry, 15(6). 10.1016/0278-5846(91)90003-J

Wong, H., & Cairns, B. E. (2019). Cannabidiol, cannabinol and their combinations act as peripheral analgesics in a rat model of myofascial pain. Arch Oral Biol, 104, 33–39. 10.1016/j.archoralbio.2019.05.028

Xiong, W., Cui, T., Cheng, K., Yang, F., Chen, S. R., Willenbring, D.,…Zhang, L. (2012). Cannabinoids suppress inflammatory and neuropathic pain by targeting alpha3 glycine receptors. J Exp Med, 209(6), 1121–1134. 10.1084/jem.20120242

Zuke, J. T., Rice, M., Rudlong, J., Paquin, T., Russo, E., & Burman, M. A. (2019). The Effects of Acute Neonatal Pain on Expression of Corticotropin-Releasing Hormone and Juvenile Anxiety in a Rodent Model. eNeuro, 6(6). 10.1523/ENEURO.0162-19.2019

Zuluaga, M. J., Agrati, D., Athaide, V., Ferreira, A., & Uriarte, N. (2023). Fear response of rat pups to a non-aversive social stimulus: Evidence for the involvement of memory processes. Dev Psychobiol, 65(7), e22417. 10.1002/dev.22417

